# Analysis of hepatic lentiviral vector transduction; implications for preclinical studies and clinical gene therapy protocols

**DOI:** 10.1101/2024.08.20.608805

**Authors:** Peirong Hu, Yajing Hao, Wei Tang, Graham H. Diering, Fei Zou, Tal Kafri

**Affiliations:** Gene Therapy Center, University of North Carolina at Chapel Hill, 27599 Chapel Hill, North Carolina, USA; Department of Biostatistics, University of North Carolina at Chapel Hill, 27599 Chapel Hill, North Carolina, USA; Department of Cell Biology and Physiology and UNC Neuroscience Center, University of North Carolina at Chapel Hill, 27599 Chapel Hill, North Carolina, USA; Carolina Institute for developmental disabilities, 27510 Carrboro, North Carolina; Department of Genetics, University of North Carolina at Chapel Hill, 27599 Chapel Hill, North Carolina, USA; Department of Microbiology and Immunology, University of North Carolina at Chapel Hill, 27599 Chapel Hill, North Carolina; Lineberger Comprehensive Cancer Center, 27599 Chapel Hill, North Carolina

## Abstract

Lentiviral vector-transduced T-cells were approved by the FDA as gene therapy anti-cancer medications. Little is known about the host genetic variation effects on the safety and efficacy of the lentiviral vector gene delivery system. To narrow this knowledge-gap, we characterized hepatic gene delivery by lentiviral vectors across the Collaborative Cross (CC) mouse genetic reference population. For 24 weeks, we periodically measured hepatic luciferase expression from lentiviral vectors in 41 CC mouse strains. Hepatic and splenic vector copy numbers were determined. We report that CC mouse strains showed highly diverse outcomes following lentiviral gene delivery. For the first time, moderate correlation between mouse strain-specific sleeping patterns and transduction efficiency was observed. We associated two quantitative trait loci (QTLs) with intra-strain variations in transduction phenotypes, which mechanistically relates to the phenomenon of metastable epialleles. An additional QTL was associated with the kinetics of hepatic transgene expression. Genes comprised in the above QTLs are potential targets to personalize gene therapy protocols. Importantly, we identified two mouse strains that open new directions in characterizing continuous viral vector silencing and HIV latency. Our findings suggest that wide-range patient-specific outcomes of viral vector-based gene therapy should be expected. Thus, novel escalating dose-based clinical protocols should be considered.

## Introduction

The first FDA approval of a gene therapy product as a cancer medication ^1^ has formally opened a new era in medicine. The successful gene therapy clinical trials that lead to the aforementioned FDA policy were premised on several lentiviral and gamma-retroviral vectors-based clinical trials ^2,3^. More recently, success in employing adeno associated virus (AAV)-based vectors to treat incurable genetic diseases further underscored the therapeutic potential of gene therapy. ^4,5^ However, due to safety concerns, the spectrum of gene therapy-treated disorders primarily includes fatal or incurable genetic and acquired diseases.

To fully realize the therapeutic potential of gene therapy, it is essential to minimize patient to patient variations in safety and efficacy of gene therapy protocols. This requirement cannot be met by current preclinical studies, in which the safety and efficacy of therapeutic viral vectors are mostly evaluated in a large cohort of rodents with an identical genetic background, and contingent with availability, in a small number of large animals (e.g., nonhuman primates). Concerns regarding the reliability of the above preclinical research system at predicting the therapeutic outcomes of viral vector-mediated gene delivery to heterogeneous patient populations were raised by a number of murine-based preclinical gene therapy studies demonstrating strain-specific efficacy and safety.^6,7,8^ Indeed, several clinical trials employing different viral vectors reported on major adverse effects, which could not be predicted by earlier pre-clinical studies. ^9-16^

Viral vector-mediated transfer of genetic cargos to target tissues and cells is an orderly process made up of multiple sequential and sometimes overlapping steps, whose completion is required for successful gene delivery. The overall efficiency of this complex multifactorial process is largely determined by the attributes of the parental virus from which a particular viral vector was derived, and by a balance of interactions between the virion’s proteins and a plethora of host factors. These host proteins and noncoding RNAs are either required for efficient completion of each step in the viral vector transduction process (dependency factors), or serve as restriction factors, which provide intrinsic immunity and are the first defense line of the host antiviral immune system.^17-21^ In a healthy human population and in mouse strains without known genetic deficiencies, expression and function levels of host dependency and restriction factors are expected to be within a relatively narrow normal range. Thus, mild non-pathologic alterations in the function/level of the abovementioned host factors are expected to significantly affect the outcomes of viral vector-based gene delivery with minimal adverse effects.

To better understand the effects of host genetic variations on the outcome of lentiviral vector-based gene therapy protocols, we sought to screen for natural allelic combinations that affect the efficacy of hepatic gene delivery by lentiviral vectors in healthy mouse strains. This approach is premised on the ability to associate phenotypic data generated *in vivo* with specific allelic combinations unique to genetically characterized mouse strains. To this end, we characterized hepatic gene delivery by lentiviral vectors in the Collaborative Cross (CC) mouse strains panel. The CC mouse strains collection is a large panel of recombinant inbred strains, which were derived from a pool of eight genetically diverse inbred mouse strains including, A/J, C57BL/6J (named below as C57B6), 129S1/SvImJ, NOD/ShiLtJ, NZO/HILtJ, WSB/EiJ, PWK/PhJ, CAST/EiJ. Altogether, the CC mouse strains panel comprises more than 36 million single nucleotide polymorphism (SNP) sites.^22^ The highly genetically diverse CC mouse strains have been successfully used for genetic mapping ^23,24^ and the development of new disease models. ^25,26,27^ Importantly, in a CC mice-based pilot study Suwanmanee *et al.*^8^ demonstrated major strain-specific differences in hepatic lentiviral vector transduction characteristics including, the level of hepatic transgene expression and the pattern of vector biodistribution.

Here we report on a large-scale genetic study, in which 254 female mice from the commonly used C57B6 mouse strain and 41 CC strains were intraperitoneally (IP) administered with lentiviral vectors expressing the firefly luciferase cDNA under the control of a liver specific promoter (hAAT). Efficacy of gene delivery was evaluated by periodic analysis of hepatic luciferase expression. Vectors copy number (VCN) in liver and spleen tissues served as a surrogate marker of vector biodistribution. A total of 23 traits of hepatic gene delivery by lentiviral vectors were phenotypically and genetically analyzed. As previously described^8^, we observed significant heritability across these measures. We located a QTL in chromosome 1, which was associated with changes in hepatic transgene expression between week 1 and week 3 -PVA lentiviral vector administration. Four genes comprised in this QTL, *Sertad4, Irf6, Traf3ip3 and UTP25* are involved in epigenetic regulation, development and the innate immune response, and thus can be considered as candidate genes contributing to the above phenomenon. In addition, we identified two QTLs, which were associated with in-strain variability (isogenic discordant) in hepatic luciferase expression at week 3-PVA and in the ratio of hepatic to spleen vector copy number (VCN). These findings suggest that isogenic discordant in host permissiveness to completion of different steps in the viral vector transduction process is relatively prevalent and genetically regulated.

In total, our results highlight the impact of genetic variation on initial responses, biodistribution and maintenance of lentivirus-based hepatic transgene delivery. The wide range of strain-specific permissiveness to hepatic lentiviral vector transduction suggests that a modified more conservative approach should be considered in designing preclinical and clinical gene therapy protocols.

## Materials and Methods

### Mice

Animal protocol (IACUC 15-270.0) and all procedures were approved by the Institutional animal care and use committee of UNC.

Collaborative cross and C57BL6 mice were obtained from UNC Systems Genetics Core Facility (SGCF).

41 Collaborative Cross strains include: CC002/Unc, CC003/Unc, CC004/TauUnc, CC005/TauUnc, CC006/TauUnc, CC008/GeniUnc, CC010/GeniUnc, CC011/Unc, CC012/GeniUnc, CC013/GeniUnc, CC016/GeniUnc, CC019/TauUnc, CC021/Unc, CC024/GeniUnc, CC027/GeniUnc, CC030/GeniUnc, CC032/GeniUnc, CC033/GeniUnc, CC035/Unc, CC036/Unc, CC038/GeniUnc, CC040/TauUnc, CC041/TanUnc, CC042/GeniUnc, CC043/GeniUnc, CC044/Unc, CC046/Unc, CC053/Unc, CC055/TauUnc, CC056/GeniUnc, CC057/Unc, CC058/Unc, CC059/TauUnc, CC060/Unc, CC061/GeniUnc, CC063/Unc, CC065/Unc, CC068/TauUnc, CC070/TauUnc, CC072/TauUnc, and CC076/Unc (named below as CC###).

### Lentiviral particles production, concentration, and titration

A lentiviral vector (pTK979) carrying the firefly luciferase cDNA under the control of a liver-specific promoter (human alpha1-antitrypsin, hAAT) was constructed and used as described earlier (Bayer 2008).^28^ VSV-G pseudotyped lentiviral vector particles were packaged with the packaging cassette ΔNRF in 293T cells using three-plasmid transient transfection as described earlier.^29^ Vector titer was determined by measuring p24 capsid concentration using p24 ELISA as previously described.^30^ All vector preps were tested for replication-competent retroviruses by 3 independent assays as described earlier.^31^

### Lentiviral vector administration and in vivo imagine

A total of 25 µg p24^gag^ VSV-G pseudotyped lentiviral vectors were intraperitoneally injected into each mouse. *In vivo* expression of vector-delivered firefly luciferase in live animals was determined at weeks 1, 3, 6, 8, 14 and 24 using IVIS Lumina optical system (PerkinElmer, Waltham, MA), as described previously.^8^

### Tissue vector copy number

Genomic DNAs from liver and spleen were isolated by using the Purelink Genomic mini-DNA kit (Invitrogen). Additional RNaseA (Fermentas, Waltham, MA) digestion step was added during DNA extraction to remove excessive RNAs. Total viral copy number (VCN) was quantified by multiplex PCR on ABI7300 Realtime PCR system as described earlier.^32^

### Sleep phenotyping and behavior analysis

Mice of the following strains C57BL/6J and CC strains 011, 013, 027, 033, 036, 041, 057, 058, 060, 072 were moved to our wake/sleep behavior satellite facility maintained on a 12 h:12 h light:dark cycle (lights on 7 am to 7 pm). Individual mice were housed in 15.5 cm^2^ cages with bedding, food, and water. Before the beginning of data collection, mice were allowed to acclimate to the environment for at least two full dark cycles. No other animals were housed in the room during these experiments. Sleep and wake behavior were recorded using a noninvasive home-cage monitoring system, PiezoSleep 2.0 (Signal Solutions, Lexington, KY), as previously described ^33^. Briefly, the system uses a Piezoelectric polymer film to quantitatively assess sleep/wake cycles, total amount of sleep and quality from mechanical signals obtained from breath rate and movement. Specialized software (SleepStats, Signal Solutions, Lexington, KY) uses an algorithm to discern sleeping respiratory patterns from waking respiratory patterns. Sleep was characterized according to specific parameters in accordance with the typical respiration of a sleeping mouse. Additional parameters were set to identify wake including the absence of characteristic sleep signals and higher amplitude, irregular signals associated with volitional movements, and even subtle head movements during quiet wake. Data collected from the cage system were binned for every 1 h to generate a daily sleep trace or 12 h bins for average light- or dark-phase percent sleep and light over dark ratios were calculated. This algorithm has been validated in adult mice by using electroencephalography, electromyography, and visual evaluation ^34-37^ and utilized successfully in additional studies ^38,39^.

### Statistical and QTL Analyses

We employed 254 female mice from C57B6 and 41 Collaborative Cross (CC) strains, with 4 to 8 replicates per strain. We measured hepatic luciferase expression at week 1, 3, 6, 8, 14, and 24 as well as liver VCN and spleen VCN. We calculated the difference in hepatic luciferase expression between different weeks, the ratio of this difference to the hepatic luciferase at the earlier week, and vector specific activity (SA) as the ratio of hepatic luciferase to liver VCN. All statistical analyses were conducted using R (version 4.2.1). To identify potential outliers, hepatic luciferase expression, VCN, and specific activity were first log transformed and then their robust estimates of mean and standard deviation were calculated (per strain) by Winsorized statistical measurements.^40^ The ratio of difference to the hepatic luciferase expression at the earlier week were added one before log transformation since some ratios were negative. Observations more than 4 standard deviations from the mean were flagged as outliers, resulting in 11 outliers across 3 phenotypes. All QTL analyses were performed on the cleaned data with the outliers being removed.

To estimate the broad-sense heritability of each phenotype, we performed a one-way ANOVA where the CC strains were modeled as a categorical variable. Broad-sense heritability was defined as the percentage of the phenotype variance explained by the CC strains, and results are summarized in table 3.

We downloaded the probability matrix of 36 genotype calls (8 homozygous calls and 28 heterozygous calls) across 76,689 SNPs from https://csbio.unc.edu/CCstatus/index.py?run=FounderProbs, which were then converted into 8-state allele probability matrix. We performed QTL mapping at each SNP using the qtl2 package in R. The genome-wide significance was empirically assessed through 1000 permutations. Phenotypes with top candidate loci that passed the genome-wide significance threshold of 0.2 are listed in table 4 with their corresponding p-value, LOD scores, Bayes credible intervals, and heritability of the top SNP.

## Results

This study is based on the notion that long-term expression of therapeutic level of lentiviral vector-delivered genetic cargos without adverse effects is the hallmark of efficacious therapeutic gene therapy protocols. However, variation in the outcomes of viral vector-based gene therapy among patients, bounds the usage of gene therapy protocols to incurable human diseases.^9-14^ We assume that the phenomenon of patient specific outcomes is secondary to the host genetic background. We believe that identification of specific host genetic loci associated with the efficacy and safety of lentiviral vector gene delivery would improve our ability to predict the outcomes of therapeutic gene delivery and would pave the way to the development of personalized gene therapy protocols. Our approach of achieving this goal, is premised on associating quantifiable phenotypes of lentiviral vector-mediated gene delivery with the genetic background of 41 genetically diverse CC mouse strains. This strategy was supported by an earlier pilot study demonstrating strong host genetic effects on the efficacy of lentiviral vector hepatic gene delivery in 12 CC mouse strains.^8^ Vector copy number (VCN) in target organs, and vector specific activity (transgene expression level per vector genome) determine long-term transgene expression level. Two intrinsic host mechanisms negatively affect long term transgene expression from lentiviral vectors. Epigenetic silencing of the transgene expression cassette would result in overall reduction in transgene expression without affecting VCN. Whilst transgene induced cytotoxicity or cellular immune response directed to vector-transduced cells would reduce VCN and overall transgene expression levels. Note that relatively rapid uncoating of HIV-1 based vectors ^41^ and the lack of earlier exposure of CC mice to HIV antigens render the possibility that cellular immune response to HIV-1 antigens will contribute to decline in transgene expression in vector transduced mice less likely, however not completely improbable.^42^ Accordingly, the study presented here aimed at identifying host genetic loci associated with the above parameters that affect long-term outcomes of lentiviral vector transduction. The study comprised two parts. A data collection part in which, we periodically quantified hepatic transduction efficiency and analyzed more than 23 traits (Tables 1-2 and Figures 1-14) of lentiviral vector-mediated hepatic gene delivery in multiple CC mouse strains. Based on these findings and on the well characterized genetic background of CC mouse strains, in the second part of this study, we identified quantitative trait loci (QTLs), which were linked by association to strain-specific characteristics of hepatic gene delivery by lentiviral vectors.

**Figure 1.**
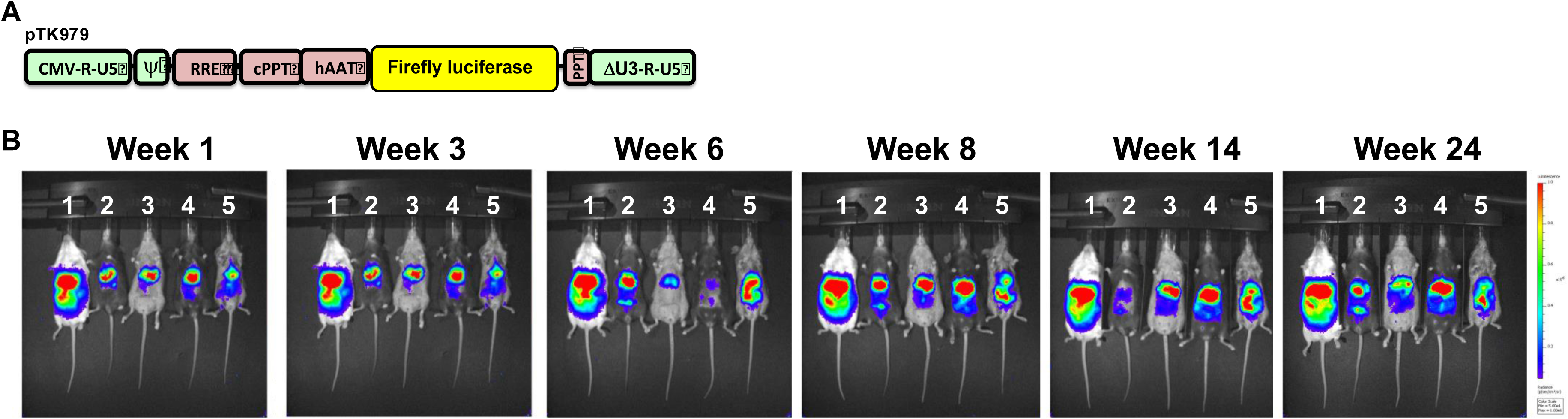
Hepatic transduction by lentiviral vectors in Collaborative Cross (CC) mice. A. Depiction of the lentiviral vector pTK979, from which the firefly luciferase is expressed under the control of the liver specific promoter hAAT. The Vector’s 5’and 3’ LTRs, packaging signal (ψ), rev response element (RRE), the central and the 3’ polypurine tract (cPPT and PPT, respectively) are shown. B. In vivo imaging of firefly luciferase expression in a sample of 5 CC mice at weeks 1, 3, 6, 8, 14 and 24-post vector administration (PVA) following intraperitoneal injection with lentiviral vectors expressing the firefly luciferase under the control of a liver specific promoter (hAAT). The mice were imaged with the IVIS *in vivo* imaging system.

**Table 1.**
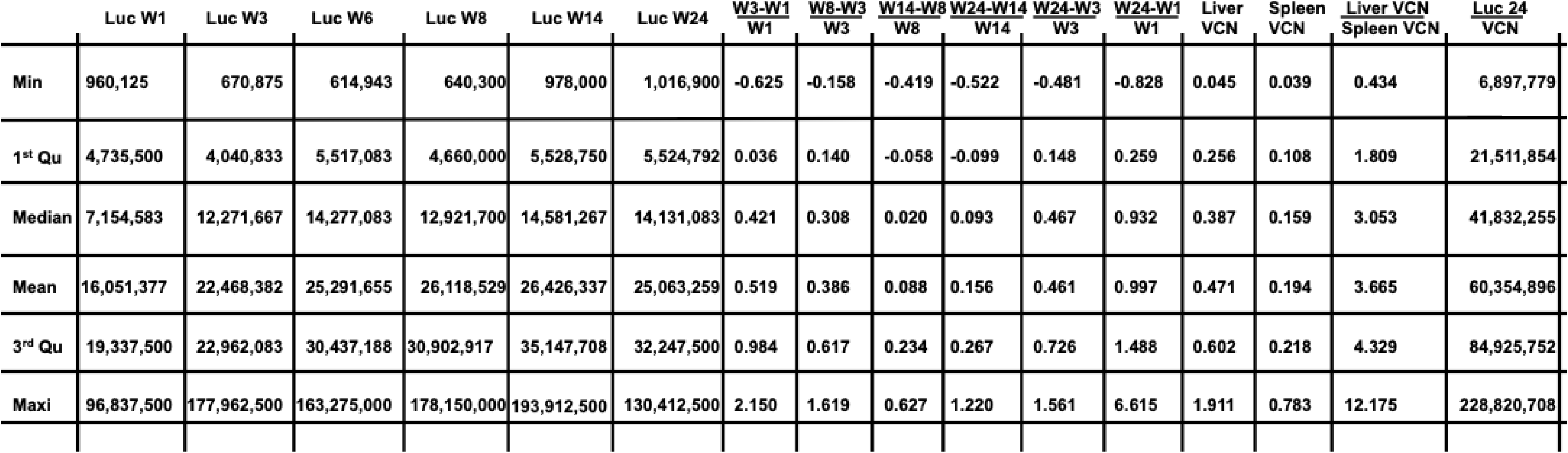
Summary statistics for the mean of each phenotype across C57B6 and 42 Collaborative Cross (CC) mouse strains. The table outlines the minima (lowest value), first quantile (25th percentile), median, mean, third quantile (75th percentile), and maxima (highest value) of the calculated means across all mouse strains.

**Table 2.** Summary statistics for the coefficient of variation (CV) of each phenotype across C57B6 and 42 Collaborative Cross (CC) mouse strains. The table outlines the minima (lowest value), first quantile (25th percentile), median, mean, third quantile (75th percentile), and maxima (highest value) of the calculated CV across all mouse strains. CV = Standard Deviation / Mean * 100%.

|  | Luc W1 | Luc W3 | Luc W24 | Liver VCN | Spleen VCN | <u>Liver VCN</u><br><u>Spleen VCN</u> | <u>Luc 24</u><br><u>VCN</u> |
| --- | --- | --- | --- | --- | --- | --- | --- |
| Min | 20.77 | 35.63 | 18.51 | 19.33 | 14.81 | 4.38 | 20.89 |
| 1<sup>st</sup> Qu | 46.87 | 52.04 | 50.02 | 32.54 | 27.44 | 40.26 | 50.43 |
| Median | 57.68 | 61.48 | 64.75 | 45.54 | 39.63 | 51.26 | 65.33 |
| Mean | 70.81 | 72.54 | 72.13 | 49.01 | 43.66 | 54.26 | 71.02 |
| 3<sup>rd</sup> Qu | 98.57 | 96.20 | 90.83 | 64.62 | 51.40 | 63.37 | 79.82 |
| Maxi | 155.21 | 184.08 | 159.87 | 95.44 | 160.22 | 135.20 | 165.26 |

**Table 3.** Phenotypes global heritability. For each phenotype that has replicated samples within each CC line, we perform one-way ANOVA analysis where the CC lines are modeled as a categorical variable. The corresponding p-value in testing the CC effect and the global heritability which is calculated as the percentage of the phenotype variance explained by the CC lines are summarized.

| Phenotype | Heritability % | p-value |
| --- | --- | --- |
| Liver Luciferase Expression Week 1 | 44.94 | <0.0001 |
| Liver Luciferase Expression Week 3 | 49.20 | <0.0001 |
| Liver Luciferase Expression Week 6 | 43.19 | <0.0001 |
| Liver Luciferase Expression Week 8 | 49.51 | <0.0001 |
| Liver Luciferase Expression Week 14 | 50.28 | <0.0001 |
| Liver Luciferase Expression Week 24 | 45.45 | <0.0001 |
| Liver Luciferase Expression Week 3 - Week 1 / Week 1 | 35.60 | <0.0001 |
| Liver Luciferase Expression Week 8 - Week 3 / Week 3 | 23.01 | 0.0370 |
| Liver Luciferase Expression Week 14 - Week 8 / Week 8 | 14.79 | 0.6819 |
| Liver Luciferase Expression Week 24 - Week 14 / Week 14 | 15.79 | 0.6130 |
| Liver Luciferase Expression Week 24 - Week 3 / Week 3 | 18.92 | 0.3018 |
| Liver Luciferase Expression Week 24 - Week 1 / Week 1 | 43.44 | <0.0001 |
| Liver VCN | 59.02 | <0.0001 |
| Spleen VCN | 53.77 | <0.0001 |
| Liver VCN / Spleen VCN | 54.28 | <0.0001 |
| Liver Luciferase expression week 24 / Liver VCN | 49.03 | <0.0001 |

**Table 4.** QTL analysis of hepatic lentiviral vector traits. For each phenotype, QTL analysis is performed, and the genome wide significance is estimated based on the result from total 1000 permutations. Phenotypes with top candidate loci that passed the genome wide significance at 0.2 are summarized. The chromosome and position (Mb) refer the estimated QTL location followed by the lower and upper bounds of the 95% Bayes credible interval and the wide of the QTL interval. The reported heritability of each QTL corresponds to the estimated heritability of the top SNP associated with the reported QTL. The numbers of genes (coding and non-protein coding) and pseudogenes located within the 95% Bayes credible interval are also reported for the three QTLs that pass the 0.05 genome wide significance level. * For each outcome, if QTL analysis is performed not on the individual measurements, but on the mean or CV of each CC line, the corresponding phenotype is referred as phenotype mean or phenotype CV, respectively.

| Phenotype * | Sig | p-value | LOD | Chr | Position (Mb) | Lower Bound (Mb) | Upper Bound (Mb) | Interval Width (Mb) | Heritability (%) | # of Genes | # of Pseudogenes |
| --- | --- | --- | --- | --- | --- | --- | --- | --- | --- | --- | --- |
| CV. Liver VCN /Spleen VCN | 0.05 | 0.008 | 9.47 | 4 | 135.41 | 134.82 | 136.43 | 1.61 | 65.48 | 57 | 11 |
| CV. Liver Luc Week-3 | 0.05 | 0.030 | 8.92 | 7 | 50.19 | 40.97 | 50.25 | 9.28 | 63.26 | 372 | 134 |
| Mean. <u>Luc Week-3 - Luci Week-1</u><br>Luci Week-1 | 0.05 | 0.040 | 8.37 | 1 | 192.69 | 192.36 | 193.71 | 1.35 | 60.92 | 34 | 4 |
| CV. Luc Week-24 / Liver VCN | 0.10 | 0.064 | 9.72 | 8 | 99.03 | 99.03 | 100.59 | 1.56 | 66.45 |  |  |
| Mean. Luci Week-1 mean | 0.10 | 0.081 | 11.27 | 2 | 154.29 | 152.95 | 156.38 | 3.43 | 71.79 |  |  |
| Mean. Liver-VCN / Spleen-VCN | 0.10 | 0.091 | 9.86 | 4 | 19.80 | 14.75 | 19.80 | 5.05 | 66.95 |  |  |
| Mean. Luci Week-6 | 0.20 | 0.143 | 10.12 | 1 | 37.56 | 37.54 | 41.37 | 3.83 | 67.91 |  |  |
| Mean. <u>Luc Week-14 - Luci Week-8</u><br>Luci Week-8 | 0.20 | 0.159 | 7.34 | 8 | 36.09 | 31.11 | 61.36 | 30.25 | 56.16 |  |  |
| CV. Spleen VCN | 0.20 | 0.166 | 11.90 | 19 | 27.49 | 24.77 | 28.06 | 3.29 | 73.72 |  |  |

### Broad range of lentiviral vector-mediated hepatic luciferase expression among 41 collaborative cross mouse strains

We sought to characterize strain-specific traits affecting the abovementioned aspects of lentiviral vector mediated hepatic gene delivery. To this end, VSV-G pseudotyped lentiviral vectors carrying the firefly luciferase cDNA under the control of a liver specific promoter (hAAT) were intraperitoneally administered to 254 females of C57B6 and 41 CC mouse strains. Hepatic luciferase expression was quantified periodically for 24 weeks (Figures 1-4).

As shown in figures 2-4 and table 1, we observed a broad range (up to 200-fold difference) of hepatic transgene (firefly luciferase) expression levels among the CC mouse strains. The means of hepatic luciferase expression among all CC mouse strains at week-1 (first measurement) and week-24 (last measurement) were, 1.61 × 10^7^ and 2.51 × 10^7^ photon/second, respectively. Strains CC061 and CC057 exhibited the highest and lowest means of hepatic luciferase expression at week-1 and week 24-PVA (9.68 × 10^7^ and 9.60 × 10^5^ photon/second at week 1 and 1.30 × 10^8^ and 1.02 × 10^6^ photon/second at week 24-PVA, respectively).

**Figure 2.**
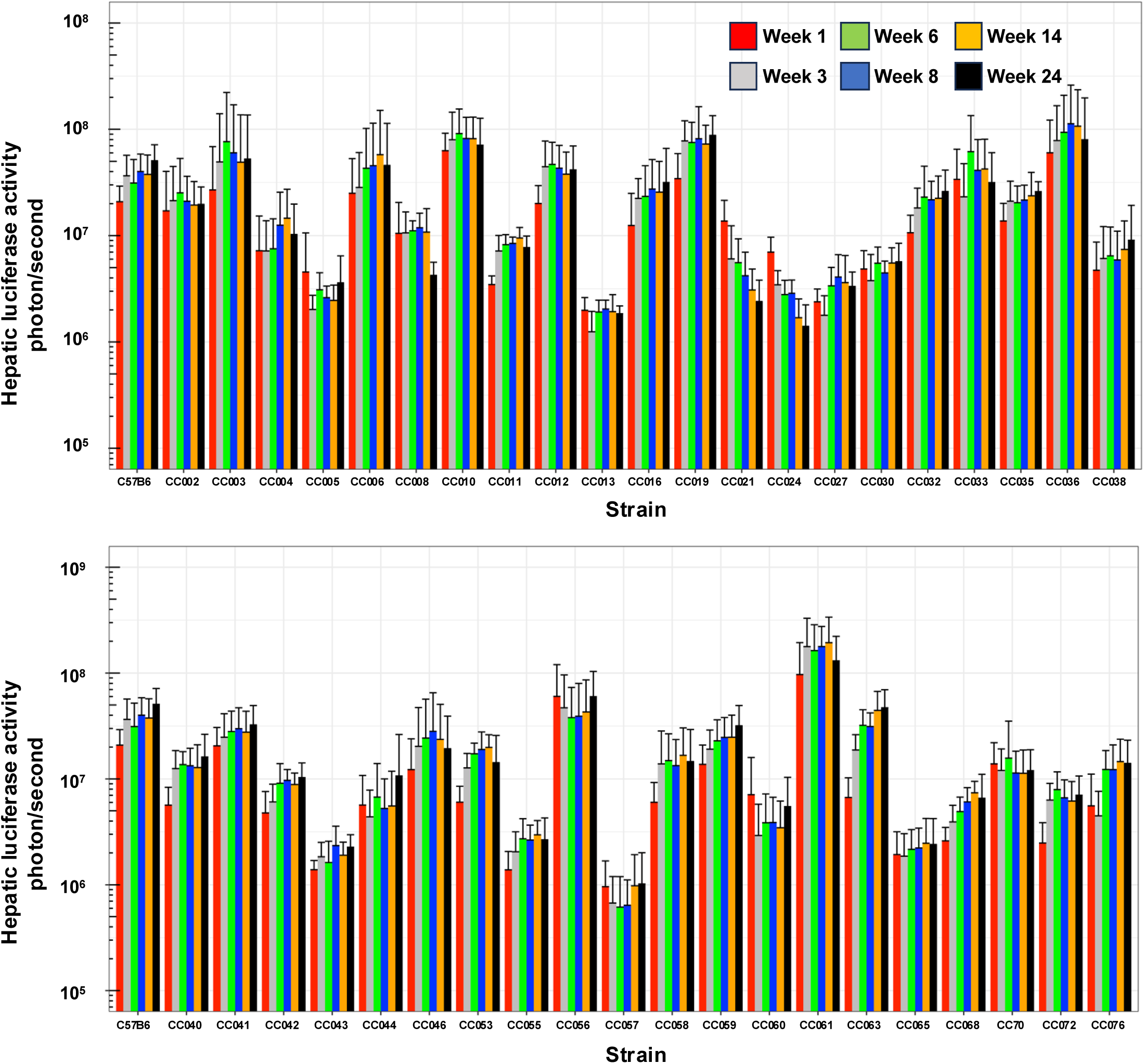
The kinetics of hepatic transgene (firefly luciferase) expression following intraperitoneal administration of lentiviral vectors. Bar graph showing levels of hepatic luciferase expression in C57B6 and 41 cc mouse strains between weeks 1 and 24-PVA. To characterize the effect of host genetic variations on hepatic gene delivery by lentiviral vectors female mice from C57B6 and 41 CC mouse strains were intraperitoneally injected with VSV-G pseudotyped lentiviral vectors (pTK979). Luciferase expression in mouse livers was periodically determined at 1,3,6,8,14 and 24 weeks-PVA by the IVIS imagine system.

**Figure 3.**
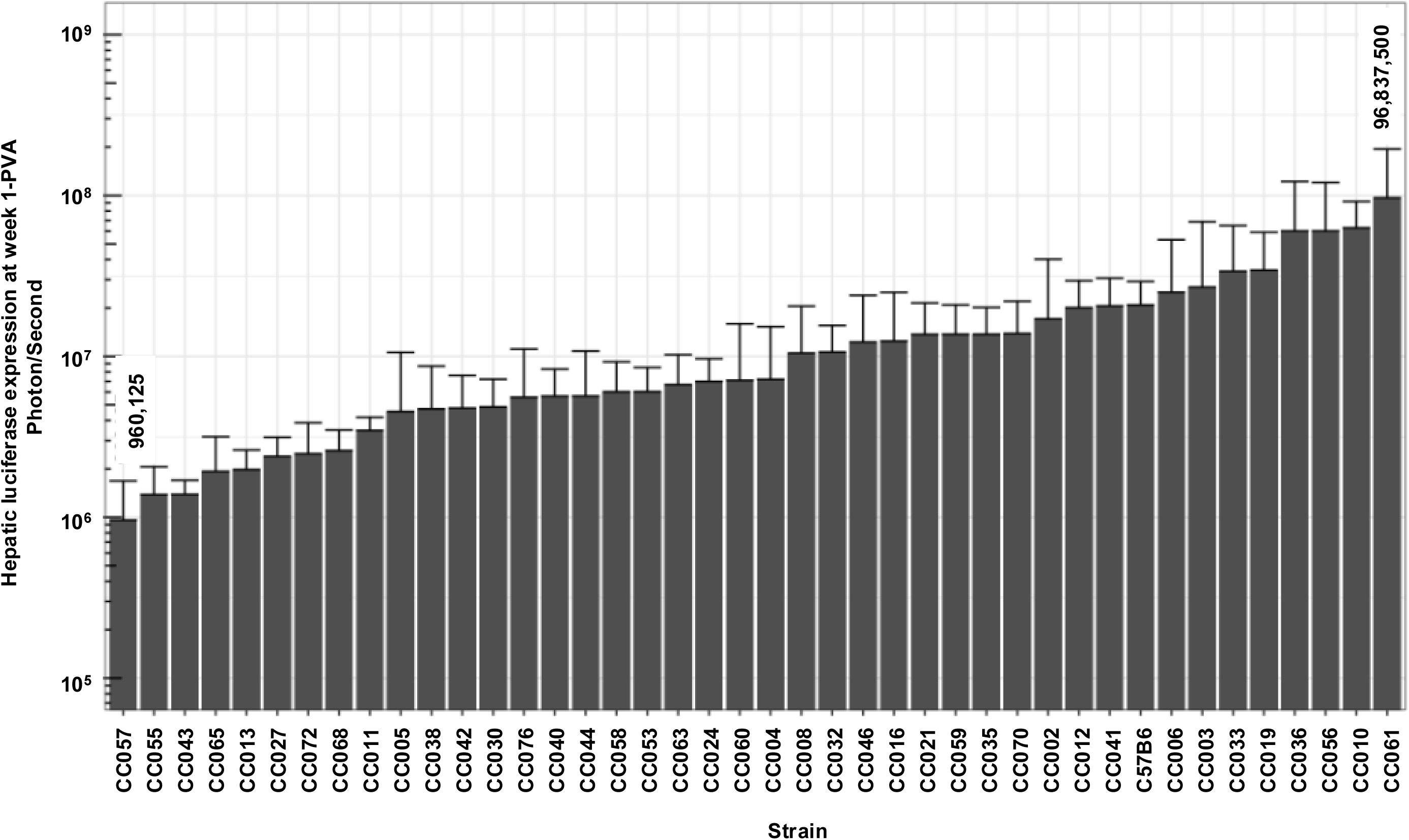
Hepatic luciferase expression at week 1-PVA in CC mouse strains. Bar graph showing the mean hepatic luciferase expression (photon/second) at week 1-PVA. The identity of the relevant CC mouse strains is shown in the bottom of the graph. The level of the lowest and highest means of hepatic luciferase expression (960,125 photon/second in CC057 and 96,837,500 in CC061, respectively) are shown.

**Figure 4.**
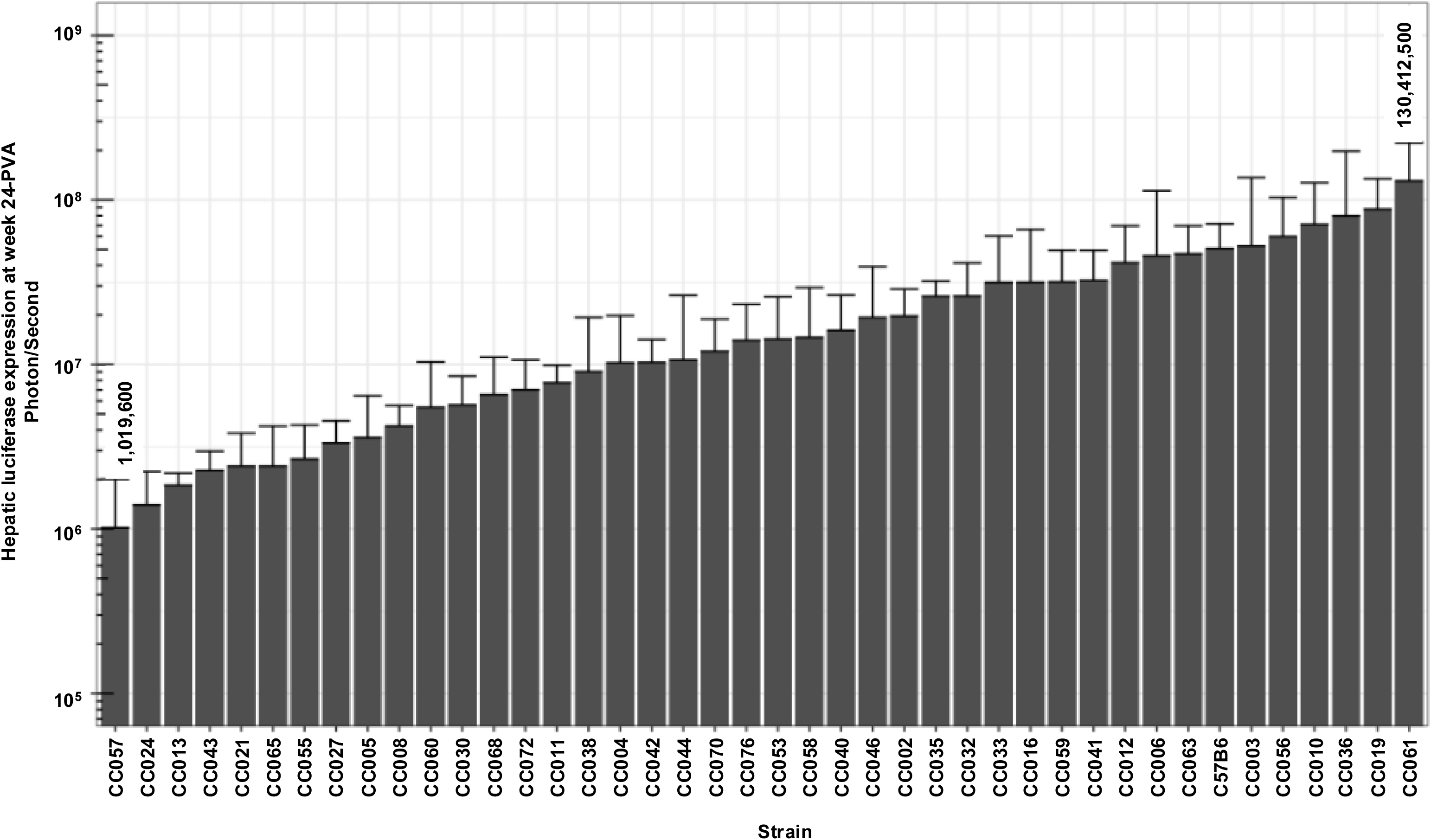
Hepatic luciferase expression at week 24-PVA in CC mouse strains. Bar graph showing the mean hepatic luciferase expression (photon/second) at week 24-PVA. The identity of the relevant CC mouse strains is shown in the bottom of the graph. The level of the lowest and highest means of hepatic luciferase expression (1,016,900 photon/second in CC057 and 130,412,500 in CC061, respectively) are shown.

As shown in figures 2-4, the means of luciferase expression levels at week 24-PVA in 15 of the 42-studied strains (35.7%) were lower than 10^7^ photon/second. In additional 25 mouse strains (59.5%) the means of luciferase expression levels at week 24-PVA were higher than 10^7^ photon/second. These data underscore the high variability in hepatic gene delivery by lentiviral vector among the CC mouse strains.

### Hepatic and splenic lentiviral vector-genome copy number per cell (VCN) in CC mouse strains

Delivery of reverse-transcribed lentiviral genomes to target cells (hepatocytes) nuclei culminates a multistep process. The efficiency of each of these steps directly affects the overall outcome of hepatic gene delivery. VCN in transduced mouse tissues is a surrogate marker of lentiviral vector gene delivery efficiency. To further characterize the effects of the host genetic background on lentiviral vector-mediated hepatic gene delivery, we employed qPCR-analysis to determine hepatic and splenic VCN in all vector transduced mice. As shown in figure 5 and table 1, we observed more than 40-fold difference between the highest and the lowest means of hepatic VCN in C57B6 (highest VCN 1.911) and CC024 (lowest VCN 0.045), respectively. The mean hepatic VCN of all studied mouse strains was 0.47 vector genome per cell. The overall mean VCN in all splenic tissues of all mouse strains, (0.19 vector genome per cell) was lower than that observed in hepatic tissues (Figure 6 and Table 1). The lowest and highest means of splenic VCN (0.04 and 0.78 respectively) were exhibited by strains CC055 and CC056, respectively. Note that these mouse strains (CC055 and CC056) are different from their counterparts that exhibited the lowest and highest hepatic VCN (CC024 and C57B6, respectively). To better understand the mechanisms affecting VCN in mouse tissues, we determined the ratio of hepatic to splenic VCN in all studied mice. As shown in figure 7 and table 1, the lowest (0.43) and highest (12.18) mean of hepatic VCN to splenic VCN ratio (in strains CC038, and CC011, respectively) differed by more than 28-fold. Altogether, these data suggest that the level of host factors affecting CC strains VCN are both strain, and cell type specific.

**Figure 5.**
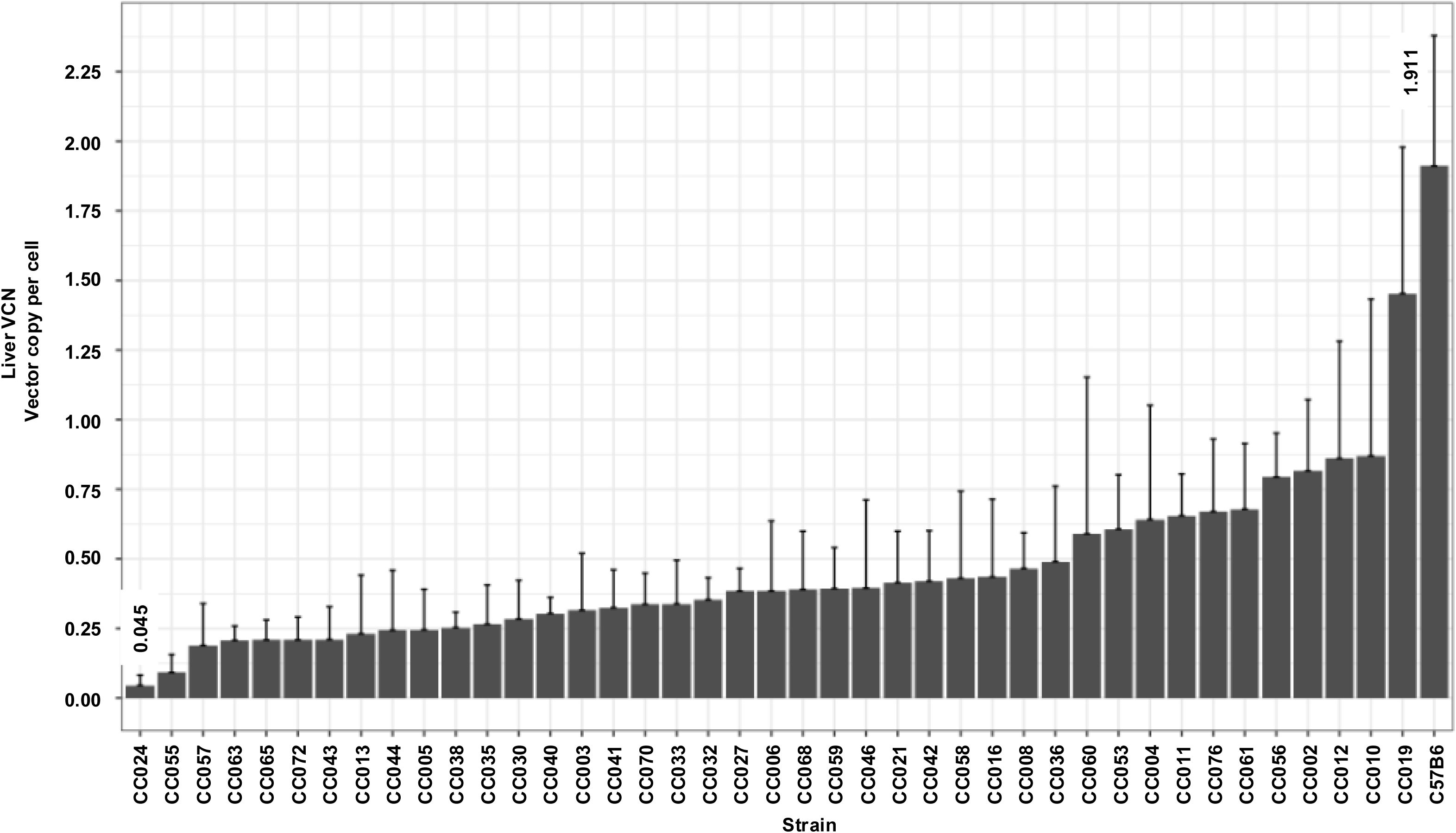
Liver VCN in CC mouse strains. Bar graph showing means of VCN in liver tissues from 41 CC mouse strains as determined by qPCR. The highest and lowest liver VCN (1.911 in C57B6 and 0.045 in CC024, respectively) are shown.

**Figure 6.**
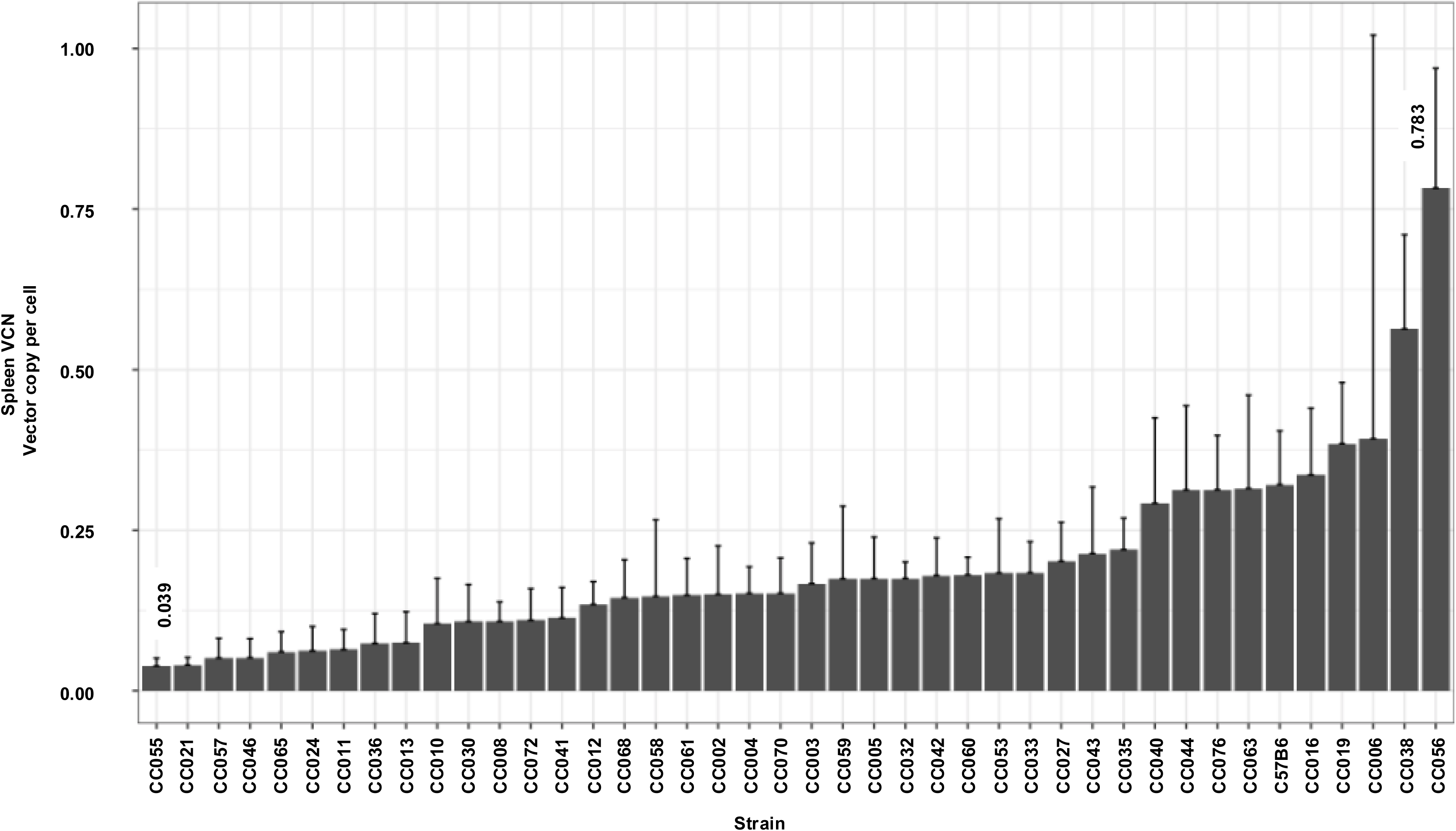
Spleen VCN in CC mouse strains. Bar graph showing means of VCN in liver tissues from 41 CC mouse strains as determined by qPCR. The highest and lowest liver VCN (0.783 in CC056 and 0.039 in CC055, respectively) are shown.

**Figure 7.**
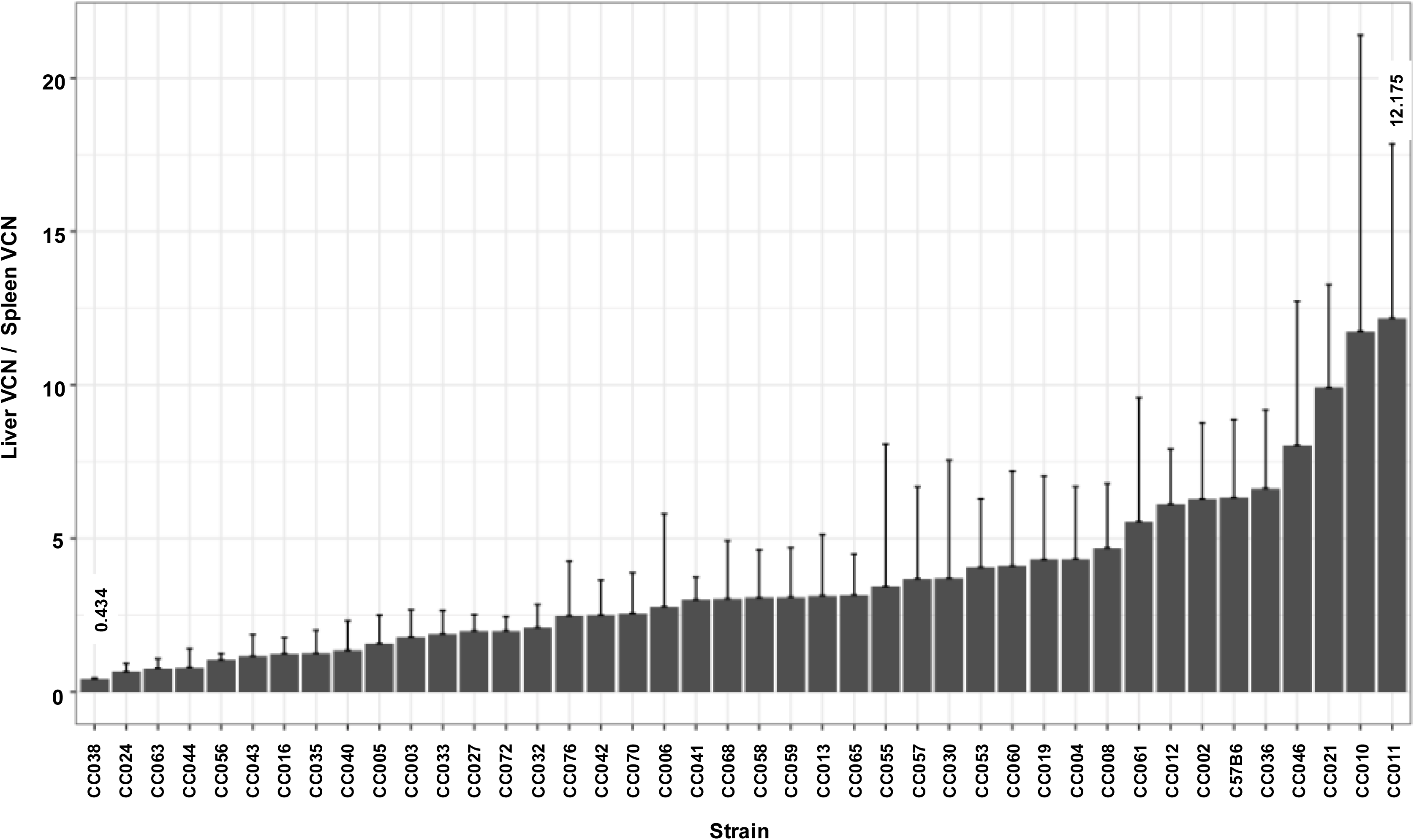
The Ratio of liver VCN to Spleen VCN. Bar graph showing the means of liver VCN to spleen VCN ratios in lentiviral vector treated CC mouse strains. VCN in the above tissues was determined by qPCR. The highest and lowest ratios in CC011 (12.175) and in CC038 (0.434), respectively are shown.

As expected, we found good correlation between the means of lentiviral vector transgene expression and the mean of hepatic VCN (Figure 8).

**Figure 8.**
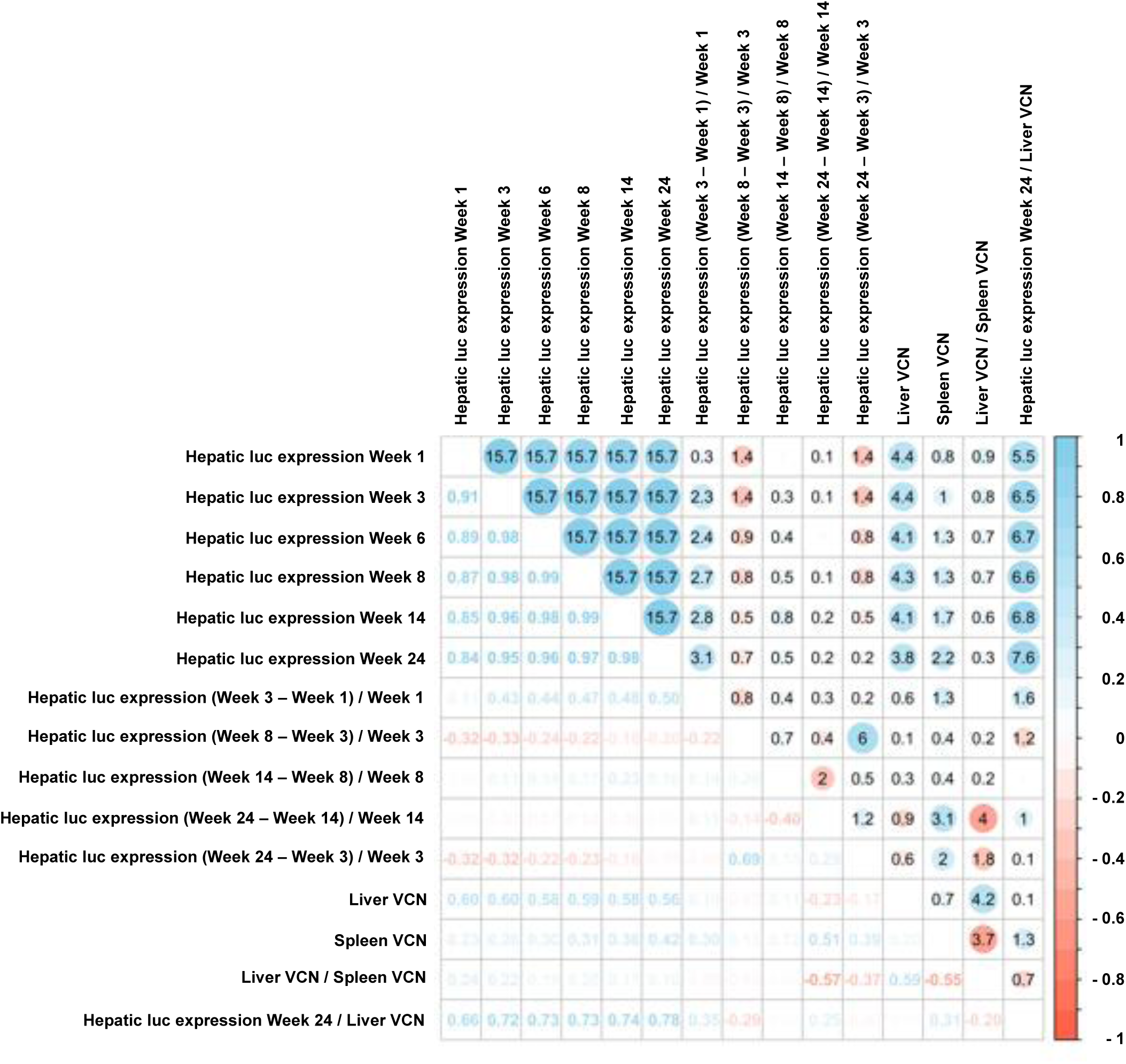
A visualization of the Spearman correlation matrix of hepatic luciferase expression and VCN. For the upper triangular correlation matrix, the areas of circles show the absolute value of corresponding correlation coefficients, and the numbers are -log10 of the p-value of tests for correlation. The positive correlations are shown in blue and negative correlations are shown in red. The lower triangular correlation matrix shows coefficients numbers with different color.

### The kinetics of hepatic luciferase expression level in CC mouse strains

The ability to maintain therapeutic levels of transgene expression is the hallmark of all successful gene replacement therapies. To study the effects of the host genetic background on the kinetics of hepatic transgene expression from lentiviral vectors, we determined the differences between hepatic luciferase expression levels at week 1-PVA and luciferase expression levels at weeks 3 and 24-PVA (Δ of W3-1 and Δ of W24-1, respectively). Since the levels of luciferase expression at week 1-PVA intrinsically affect the absolute value of the above Δs (Figure 9), we normalized the effects of W1-PVA expression levels by dividing the values of the above Δs with the values of their respective expression levels at W 1-PVA (ΔW3-1/W1 and ΔW24-W1/W1). As shown in figures 2, 10-11 and table 1, C57B6 mice and most of the CC mouse strains exhibited a stable or an increased hepatic luciferase expression throughout the study. However, two mouse strains, CC021 and CC024 showed a significant decrease in hepatic transgene expression levels between Weeks 1- and 24-PVA. Specifically, W24- W1/W1 of -0.81 and – 0.76 were calculated for CC021 and CC024, respectively. This phenomenon could have been mediated by, either specific elimination of luciferase-expressing hepatocytes or/and by transcriptional silencing of the lentiviral vector expression cassettes. We theorized that VCN in hepatic and splenic tissues in CC021 and CC024 mouse strains could imply on the role of the above two mechanisms in the decrease of hepatic transgene expression from lentiviral vectors. As shown in figures 5 and 7, hepatic tissues in CC024 mice exhibited the lowest levels of VCN, and the second lowest hepatic VCN to splenic VCN ratio. These findings support the hypothesis that elimination of luciferase-expressing hepatocytes probably contributed to the decrease in hepatic transgene expression from lentiviral vectors. Per contra, CC021 mice exhibited moderate levels of hepatic VCN and the third highest ratio of hepatic to splenic VCN. Based on these data, we suggest that transcriptional silencing of the lentiviral vector cassettes contributed to the decrease in hepatic luciferase expression between weeks 1 and 24-PVA in CC021 mice.

**Figure 9.**
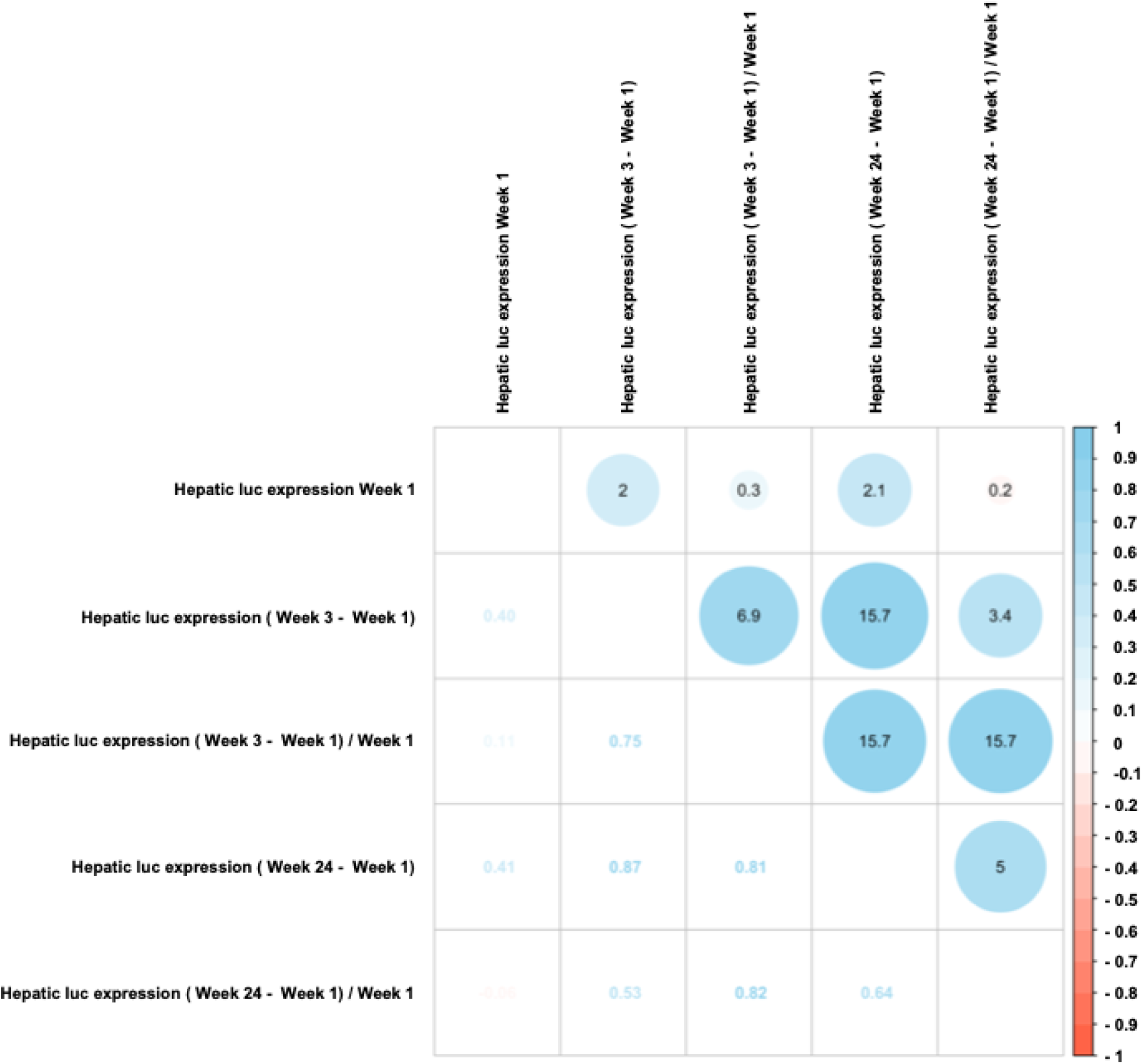
A visualization of the Spearman correlation matrix of hepatic luciferase expression at week 1-PVA and the differences in luciferase expression levels between week 1-PVA and weeks 1 and 24-PVA. For the upper triangular correlation matrix, the areas of circles show the absolute value of corresponding correlation coefficients, and the numbers are -log10 of the p-value of tests for correlation. The positive correlations are shown in blue and negative correlations are shown in red. The lower triangular correlation matrix shows coefficients numbers with different color.

### Lentiviral vector specific activity in CC mouse strains

The above findings implied that strain-specific transcriptional regulation of lentiviral vector cassettes controls hepatic transgene expression. Thus, we sought to characterize specific activity (SA) of lentiviral vectors carrying a liver specific promoter (hAAT) in the above panel of CC mouse strains. To this end, vector SA activity in each of the studied mice was calculated by dividing mouse hepatic luciferase expression level at week 24-PVA by mouse hepatic VCN. The CC mouse panel exhibited a wide range of mean lentiviral vector SA (Figure 12 and Table 1). Specifically, more than 30-fold difference in vector SA were observed between the lowest and highest means of vector SA, (6.9 × 10^6^ photon/second/VCN in mouse strain CC021 and 2.3 × 10^8^ photon/second/VCN in mouse strain CC063, respectively). These findings indicate that in addition to VCN, vector specific activity which is determined by the host transcription factors profile and the epigenetics of the vector affect the outcomes of gene delivery in a strain specific manner.

**Figure 10.**
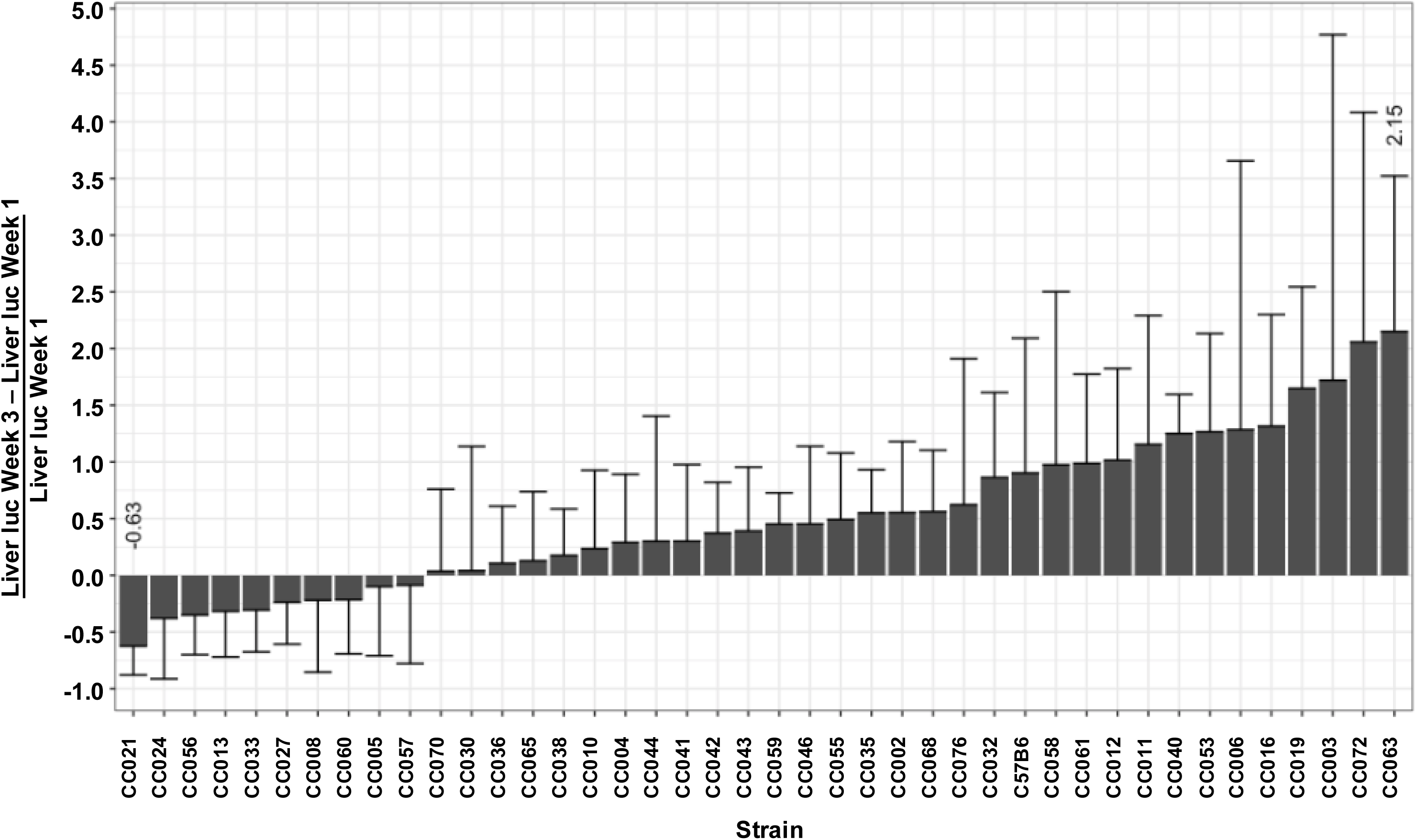
The kinetics of hepatic transgene expression between week 1 and week 3 -PVA. Bar graph showing the difference in the mean hepatic transgene expression between week 1-PVA and week 3-PVA in the abovementioned 41 CC and C57B6 mouse strain. To minimize the intrinsic effect of strain specific transgene expression at week 1-PVA on the magnitude of the difference in transgene expression at weeks 1 and 3-PVA, the calculated differences were divided by the value of transgene expression at week 1-PVA. The highest and lowest means (2.15 in CC063 and -063 in CC021, respectively) are shown.

**Figure 11.**
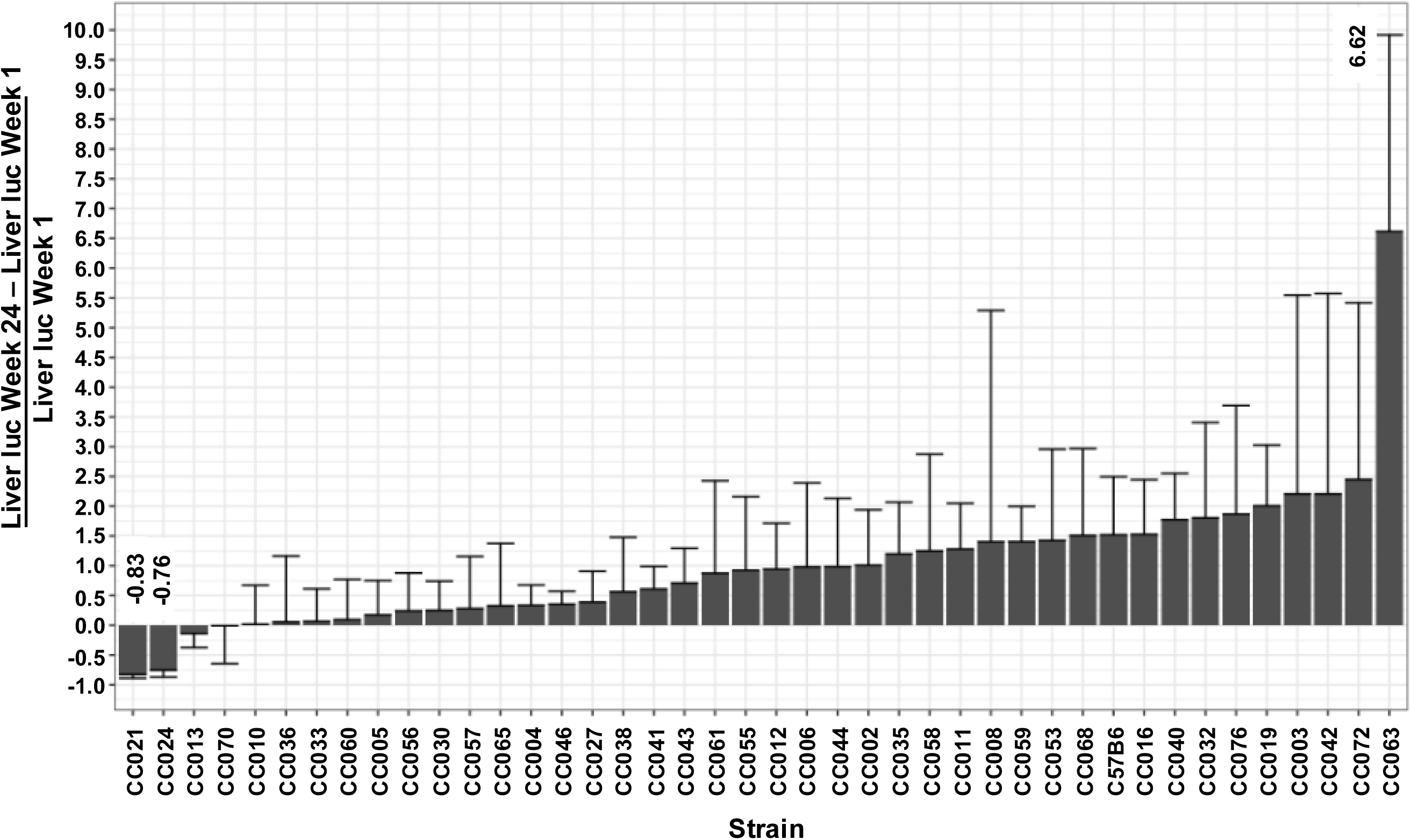
The kinetics of hepatic transgene expression between week 1 and week 24-PVA. A bar graph showing the difference in the mean hepatic transgene expression between week 1-PVA and week 24-PVA in the abovementioned 41 CC and C57B6 mouse strain. To minimize the intrinsic effect of strain specific transgene expression at week 1-PVA on the magnitude of the difference in transgene expression at weeks 1 and 24-PVA, the calculated differences were divided by the value of transgene expression at week 1-PVA. The highest and lowest means (6.62 in CC063 and -0.83 in CC021, respectively) are shown.

**Figure 12.**
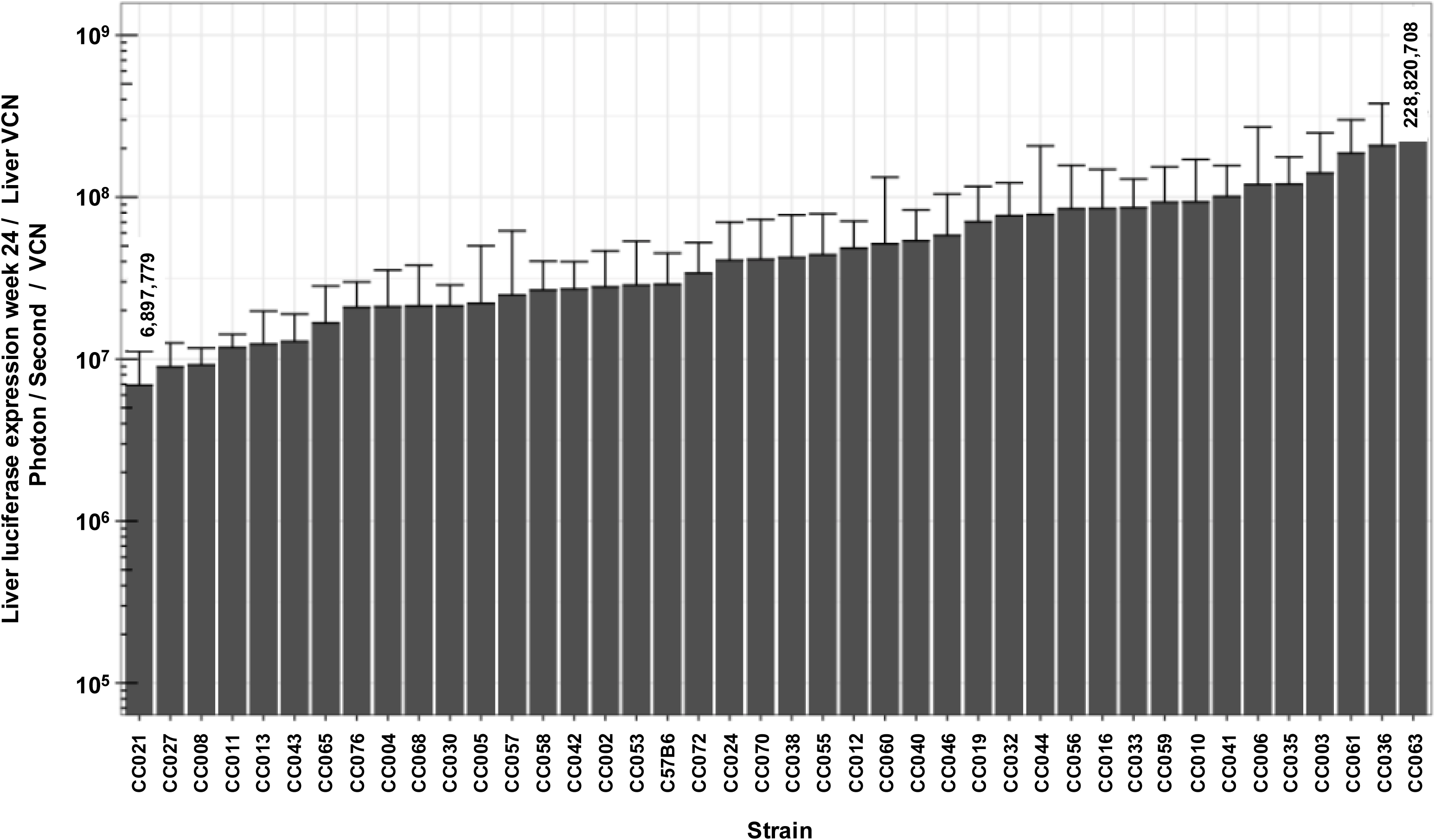
Specific activity of lentiviral vectors carrying a liver specific promoter in CC mouse strains. Bar graph showing means of vector specific activity in C57B6 and 41 CC mouse strains. To determine vector specific activity, hepatic luciferase activity at week 24-PVA was divided by liver VCN. The highest and lowest means vector specific activity (228,820,708 photon/second/vector copy per cell, in CC063 and 6,897,779 photon/second/vector copy per cell in CC021, respectively) are shown.

### Correlations between different characteristics of hepatic transduction by lentiviral vectors

To better understand the mechanisms affecting hepatic transduction efficiency by lentiviral vectors, we calculated the correlation between the above traits (Figure 8). Hepatic luciferase expression levels in weeks 1- to 14-PVA highly correlated with hepatic luciferase expression at week 24-PVA. A mild increase in Spearman correlation was observed between weeks 1-PVA/24-PVA (0.84) and weeks 3-PVA/24-PVA (0.95). As expected, the levels of hepatic VCN positively correlated with hepatic transgene expression throughout the study (R ≥ 0.56). Vector SA (hepatic luciferase expression at week 24-PVA / VCN) positively correlated with hepatic luciferase expression at weeks 1 to 14-PVA. The lowest level of Spearman correlation of vector SA was observed with week 1-PVA (0.66). A mild increase in Spearman correlation was observed at week 3-PVA (0.72) and was comparable to the Spearman correlation at weeks 6,8 and 14-PVA). Except for Spearman correlation between hepatic luciferase expression at week 24-PVA and the differences observed in hepatic luciferase expression at weeks 1 and 3-PVA (0.50), differences in hepatic luciferases expression between the different time points during the study did not correlate with hepatic luciferase expression at week 24-PVA. Altogether, these findings raise the possibility that changes in hepatic luciferase expression level and vector SA between weeks 1 and 3-PVA positively affect long-term hepatic transgene expression from lentiviral vectors.

### In-strain variabilities in characteristics of hepatic transgene expression

Intrigued by the wide range of phenotypes, amongst the different CC mouse strains, we sought to characterize intrastrain variabilities in the above studied traits. To this end, coefficient variation (CV) of the above traits was calculated (CV=standard deviation/mean) for each CC mouse strain. As shown in figures 13-14 and table 2, we observed a wide range of CVs of the studied phenotypes within the different CC mouse strains. These findings indicated that the host genetic background affect both strain and intrastrain specific phenotypes and raised the possibility that specific host factors contribute to overall intrastrain phenotypic variability. Specifically, we sought to investigate if an increased CV in a specific phenotype increases the likelihood of observing an increase in CV of other phenotypes. As shown in a heatmap representation (Figure 15) specific mouse strains exhibited increased CV in several phenotypes some of which were not directly related. These findings suggest that the host genetic background could contribute to the overall genotypic phenotypic discordance in a trait specific manner and potentially the ability to respond to environmental pressures.

**Figure 13.**
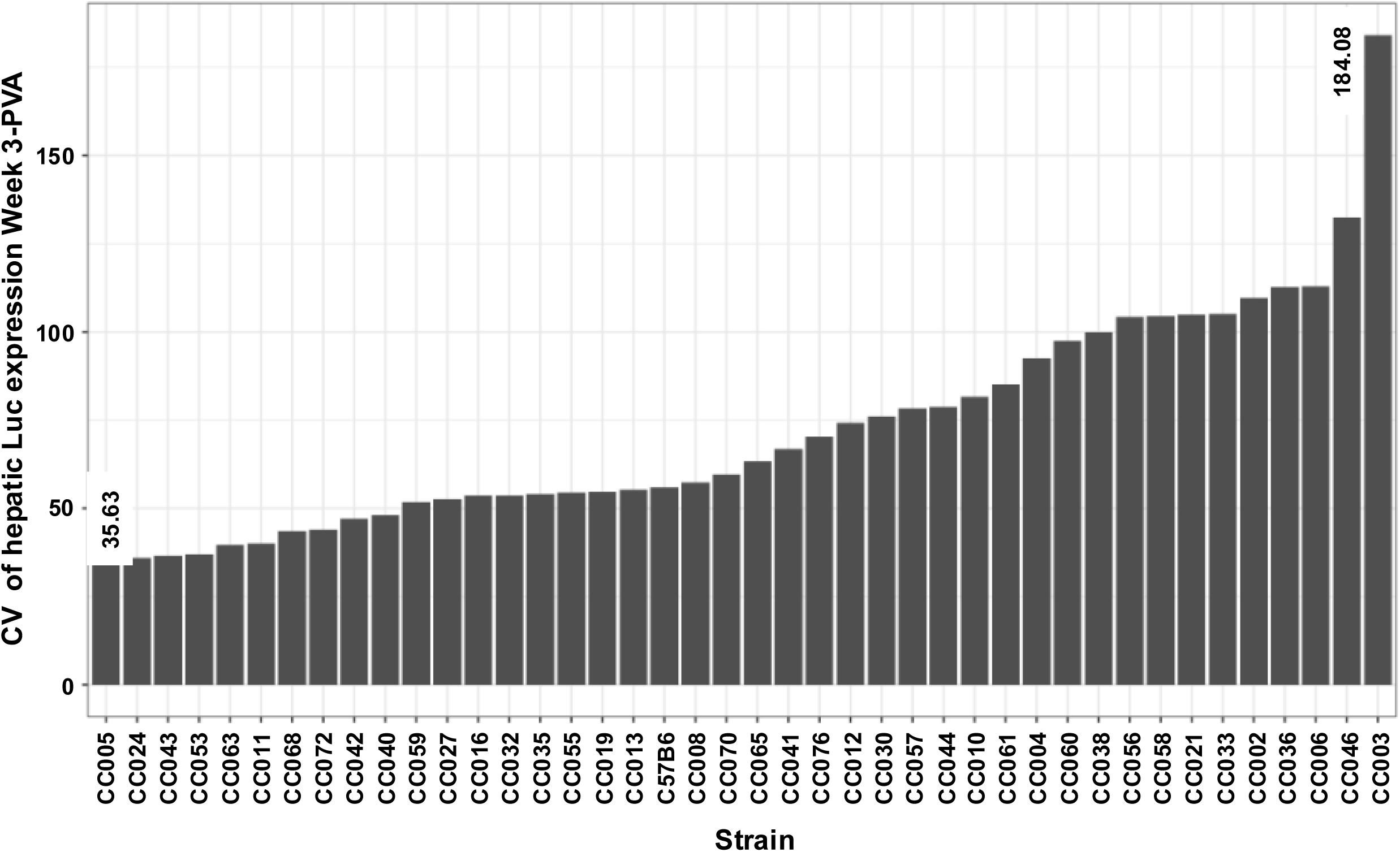
Coefficient variation (CV) of hepatic luciferase expression at week 3-PVA. Bar graph showing mouse strain specific intrastrain variation in luciferase expression at week 3-PVA. Since mean luciferase expression levels intrinsically affect standard deviation (std) of luciferase expression. Coefficient variation (std/mean) was employed to characterize strain specific effects on intrastrain variation in luciferase expression at week 3-PVA. The highest and lowest CV (184.08 in strain CC003 and 35.63 in CC005, respectively) are shown.

**Figure 14.**
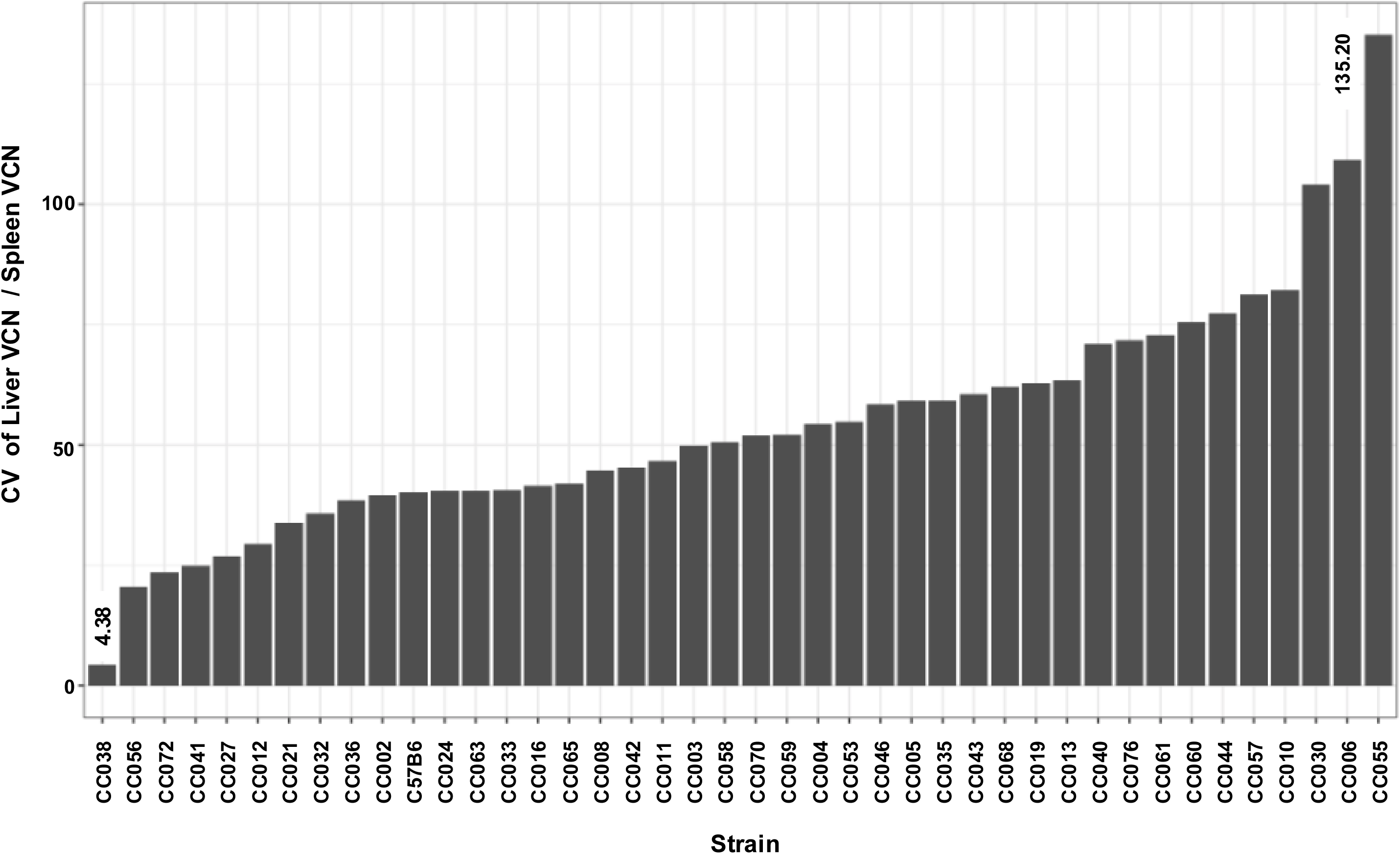
CV of liver VCN and spleen VCN ratios (liver VCN/Spleen VCN) in CC mouse strains. Bar graph showing mouse strain specific intrastrain variation in liver *VCN/spleen VCN ratios*. Since the mean of the above ratios intrinsically affects the std of the above calculated ratios within each mouse strain, coefficient variation (std/mean) was employed to characterize strain specific effects on intrastrain variation in liver VCN / Spleen VCN ratios. The highest and lowest CV (135.20 in strain CC055 and 4.38 in CC038, respectively) are shown.

**Figure 15.**
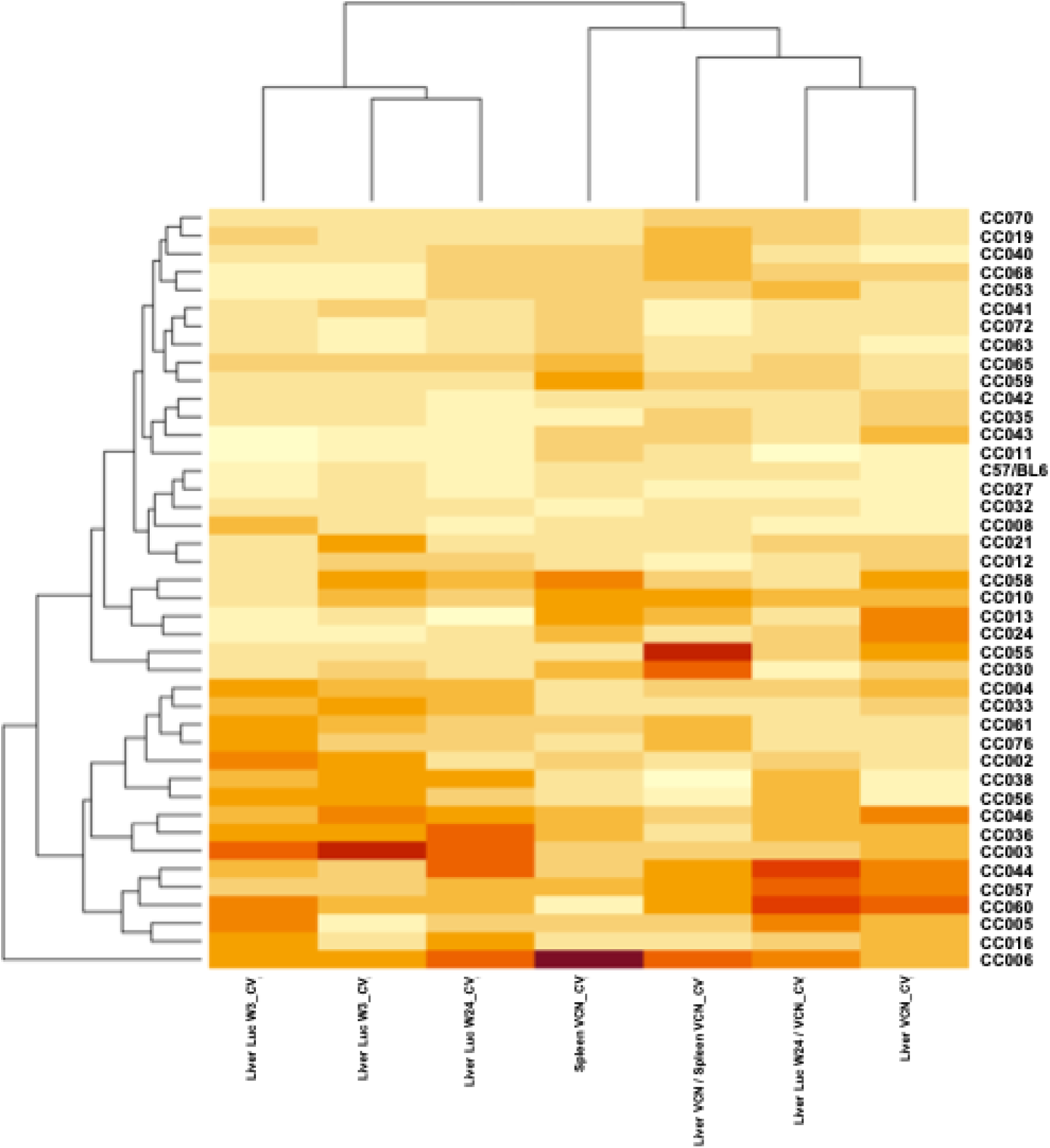
Heatmap of variables related to CV across mouse strains. Values are displayed as colors ranging from light yellow to dark red, and larger values are in darker colors. Both rows and columns are clustered using correlation distance with a dendrogram added to the left side and to the top.

### The overall effect of the host genetics on hepatic transduction by lentiviral vectors

To quantify the contribution of the host genetics to the characteristics of hepatic transduction by lentiviral vector in the CC mouse strain panel, we calculated global heritability of each phenotype that had replicates within each CC mouse strain. The corresponding p-values in testing the CC genetic effects and the global heritability are summarized in table 3. As shown, heritability of most phenotypes (12 out of 16) ranged between 43.19% to 59.02 % with p-values <0.0001. Heritability of four phenotypes which focused on the kinetics of hepatic transgene expression after week 3-PVA was low (14.79%-23.01%) with p-values 0.613-0.037. Per contra the global heritability of changes in hepatic luciferase expression between weeks 1 and 3-PVA was highly significant (p-value <0.0001) and thus, further support the notion that changes in hepatic transgene expression from lentiviral vectors are likely to occur during the first three weeks post transduction. Overall, these data confirmed our notion that the host genetic background has a major role in establishing strain-specific characteristics of lentiviral vectors transduction. However, it appeared that non-genetic factors also contributed to the above studied phenotypes.

### Mouse strain specific sleep patterns as an intrinsic environmental factor affecting hepatic transduction by lentiviral vectors

The above findings suggested that in addition to the significant host-genetic effects on hepatic lentiviral transduction, environmental factors probably contributed to the studied phenotypes. Importantly, in this study, all *in vivo* experiments were carried under a tightly controlled environment. We theorized that mouse strain-specific sleeping patterns could generate an intrinsic environment, which directly affect adult mouse gene expression. Note that daily animal care (e.g. bedding replacement, cage care) and treatments in most academic facilities takes place during the mouse daylight/sleep period, which can result in continuous sleep disturbances, whose magnitude of effects could be mouse strain specific. Furthermore, there is a possibility that aberrant maternal sleep patterns induce life-long behavioral and physiological alterations. To study the potential effects of sleep patterns on hepatic lentiviral vector transduction, we made use of sleep/awake behavior patterns, which were analyzed in a separate earlier experiment by PiezoSleep in 11 random mouse strains (C57B6 and 10 CC mouse strains). The PiezoSleep system is an automated, non-invasive home-cage monitoring system that uses vibrational sensors to measure mouse motion and breathing to score wake/sleep behavior.^34,35,38,39^ Mice are nocturnal; thus, it is expected that mice will spend most of their sleeping time during the light period (Figure 16-A). The percent of sleep time was determined hourly for 7 days in the above mouse strains (Figures 16 A-B). To quantify sleep patterns behaviors, the averages of the ratios of percent sleeping time during light (zeitgeber 1-12) and dark (zeitgeber 13-24) periods were calculated and demonstrated significant strain-specific phenotype (Figure 16-B). Next, we calculated the correlation between the light/dark sleep ratios in the above 11 mouse strains and hepatic luciferase expression at week 24-PVA which were determined in the respective mouse strains in the current study. As shown in figure 16-C, moderate yet significant correlation (p-value = 0.028 and Pearson’s correlation = 0.66) was observed between the levels of hepatic transgene expression and the ratio of female mouse sleeping time during light and dark periods. Although, highly intriguing, these findings should be further supported by using a larger number of CC mouse strains.

### Identification and characterization of QTLs associated with traits of hepatic lentiviral vector transduction

To better understand the role of the host genetic background in the process of hepatic transduction by lentiviral vectors, we sought to identify genomic loci responsible for transcribing either protein-encoding or non-protein coding RNAs that affect the above studied phenotypes. For this purpose, we performed a quantitative trait locus (QTL) analysis using qtl2 package in R. Specifically, the significance of the linkage between genetic marker (SNPs) and the quantitatively characterized lentiviral vector hepatic transduction traits was determined by the LOD score (log10 of odds). This method measures the ratio between the probability of associating strain-specific SNPs with a linkage to the studied trait to the probability of associating strain specific SNPs in the absence of linkage. The genome-wide significance was estimated based on the result from 1000 permutations.

Three trait-associated QTLs with p-values <0.05 were identified (Table 4). The above traits include coefficient variation (CV) of the ratio between liver VCN to spleen VCN, CV of hepatic luciferase expression at week 3-PVA, and the difference between luciferase expression at week 3 and 1 -PVA divided (normalized) by luciferase expression at week 1-PVA.

### QTL association with CV of the ratio between liver VCN to spleen VCN

The phenomenon of strain-dependent intrastrain variations in traits of hepatic lentiviral vector transduction (Figures 13-14 and Table 2), raised the possibility that this phenomenon is affected by the host genetic background. This concept suggests that the host genetic background determines the magnitude of isogenic phenotypic discordance of specific traits and potentially the contribution of environmental effects on this phenomenon. To test this hypothesis and to identify QTLs associated with intrastrain variations, CV of hepatic lentiviral vector transduction phenotypes were treated as traits. This approach was premised on the notion that in contrast to standard deviations (SDs), CVs are less affected by the size of the respective means. Thus, the likelihood of identifying genetic loci, which directly affect a CV of a trait is higher compared to those affect SD. Indeed, as shown in figure 17, both the means of strain luciferase expression at week 3-PVA and the means of the ratios of liver-VCN to spleen-VCN correlated better with strain SDs (R = 0.968 and 0.893, respectively) than with strain CVs (R = 0.414 and 0.192, respectively). As shown in figures 18 A-D and table 4, QTL analysis of CV of the ratios of liver-VCN means to spleen-VCN means, identified in chromosome 4 a single QTL with LOD score of 9.47 and p-value of 0.008. The QTL is positioned in Mb 135.41 and its width is 1.61 Mb. It comprises 33 protein-coding genes and 11 pseudo-genes. The percent of contribution (heritability) from the top SNP in QTL regions to the above phenotype is 65.48%. As shown in figures 18 C-D. 4 CC (CC010, CC030, CC055, CC057) mouse strains, which are composed of the 129s1/SvImJ allele in the above QTL are associated with the highest intrastrain differences (CV) in the ratio of liver VCN to spleen VCN. Aware of the role of CCCTC-binding factor (CTCF) and retroelements in the phenomenon of isogenic phenotypic discordance and genomic imprinting, ^43-46^ we mapped CTCF binding sequences in the QTL region as well. As shown in figure 19. 247 CTCF binding sites were identified in the above QTL. 7 129s1/SvImJ-unique retroelements insertions were identified including 5 SINES, an RLTR10 and a MaLr.^45^ Furthermore, the location of the above MaLr insertion in the CTCF cluster raises the possibility that in line with earlier reports interaction between retroelements and CTCF binding sequences may contribute to the increased CV of the above phenotype.^43-47^

**Figure 16.**
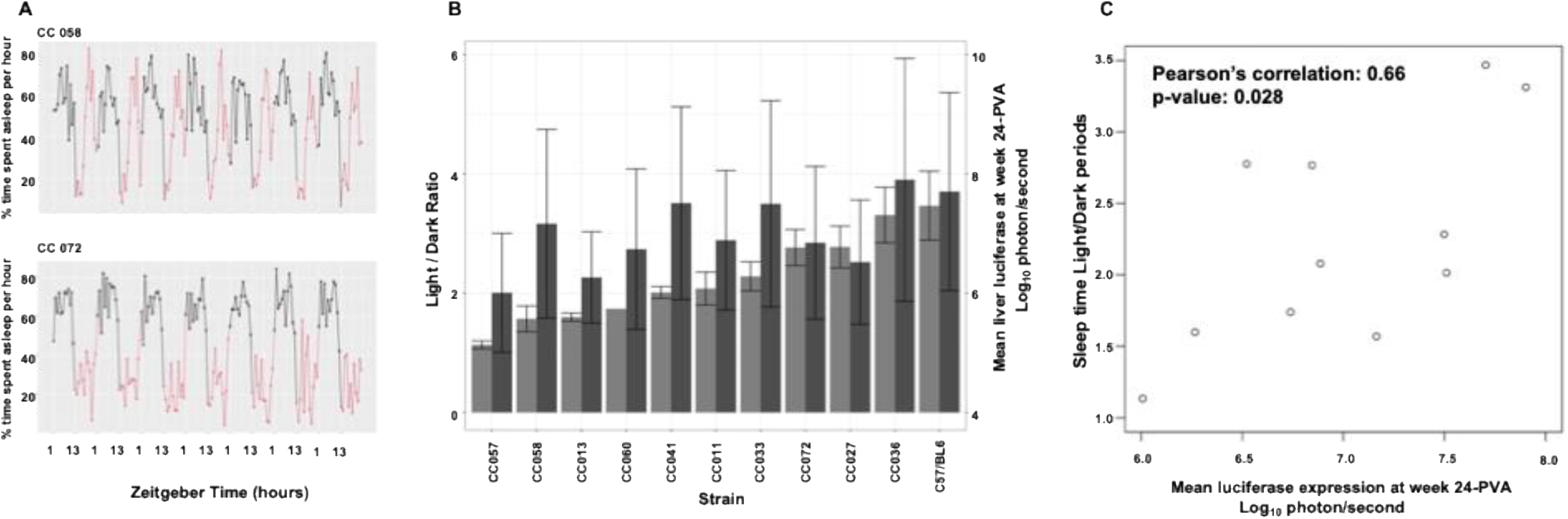
The effects of sleep patterns on hepatic transduction by lentiviral vectors. Figure 16A. The average percentage of time spent asleep per hour in 7-day sleep traces for CC058 and CC072. The x axis represents zeitgeber time in hours. Black dots indicate lights on, and red dots indicate lights off. Figure 16B. Bar plots show the average ratio of time spent asleep during light periods compared to dark periods (light grey) and the log10 of mean liver luciferase at week 24 (dark grey) for each Collaborative Cross mouse strain. The error bars represent one standard deviation, with half a standard deviation above and below the mean. We calculated the average percentage of time spent asleep during light or dark periods for each mouse, then computed the ratio of time spent during light to dark periods, and finally averaged ratios across all mice for each strain. Figure 16C. Scatter plots show the relationship between the average ratio of time spent asleep during light periods compared to dark periods and the log10 of mean liver luciferase at week 24. Pearson’s correlation = 0.6583 (p-value = 0.0277). There are 5 female mice in C57/BL6, 5 in CC011, 5 in CC013, 4 in CC027, 5 in CC033, 4 in CC036, 4 in CC041, 5 in CC057, 5 in CC058, 1 in CC060, and 4 in CC072, respectively when calculating the ratio of time spent asleep during light periods compared to dark periods. There are 8 female mice in C57/BL6, 5 in CC011, 6 in CC013, 7 in CC027, 6 in CC033, 8 in CC036, 6 in CC041, 8 in CC057, 6 in CC058, 6 in CC060, and 6 in CC072, respectively when calculating the mean liver luciferase at week 24.

**Figure 17.**
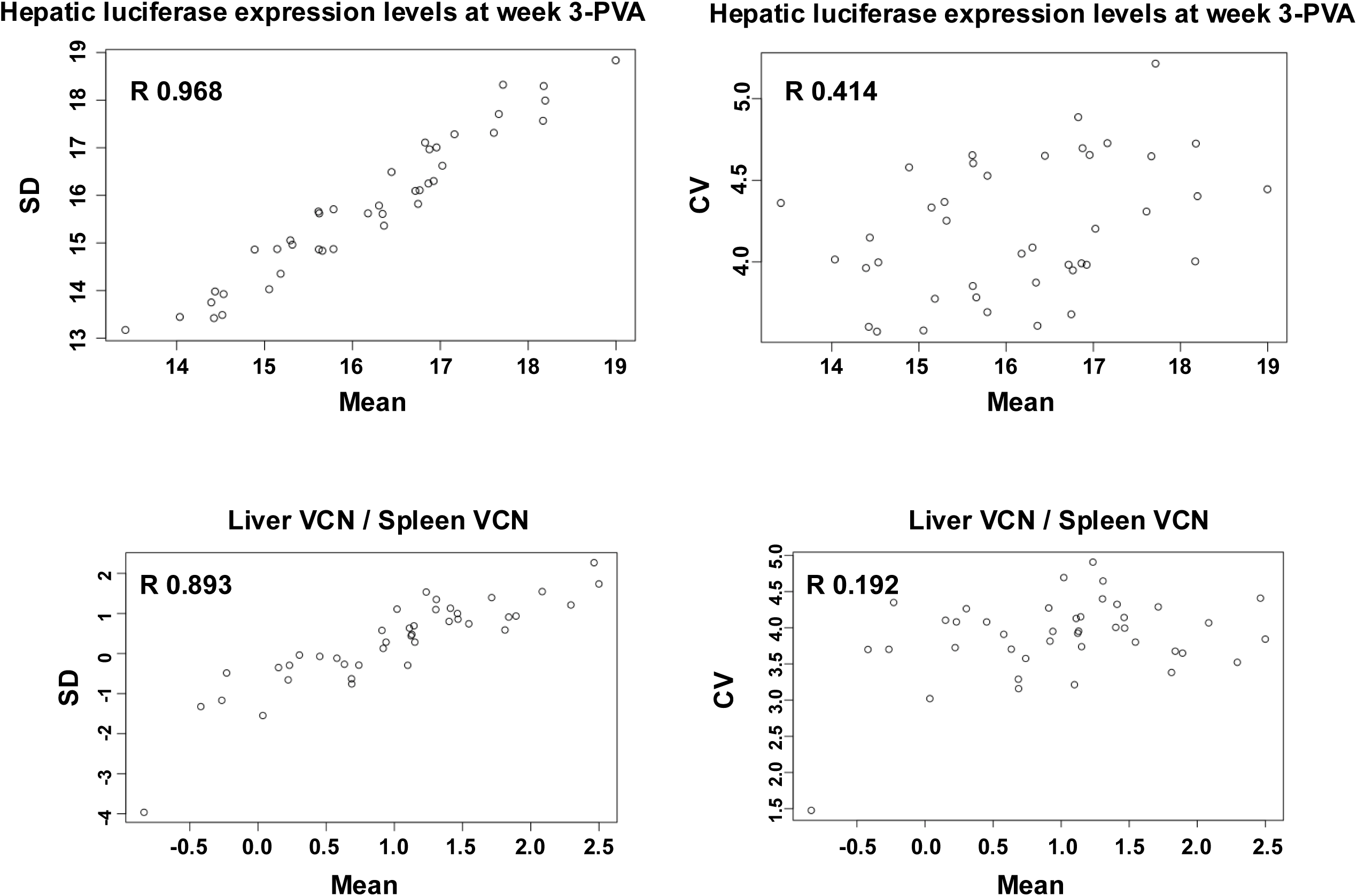
The intrinsic effects of mean luciferase expression levels on standard deviation (SD) and coefficient of variation (CV). Scatter plots show the relationship between the SDs and means of hepatic luciferase expression at week 3, CV and mean of hepatic luciferase expression at week 3, SD and mean of the ratio of liver and spleen VCN, and CV and mean of the ratio of liver and spleen VCN, respectively. The corresponding correlation coefficients are added. The correlation between SD and mean of hepatic luciferase expression at week 3 is high, and the dots fall along the line.

**Figure 18.**
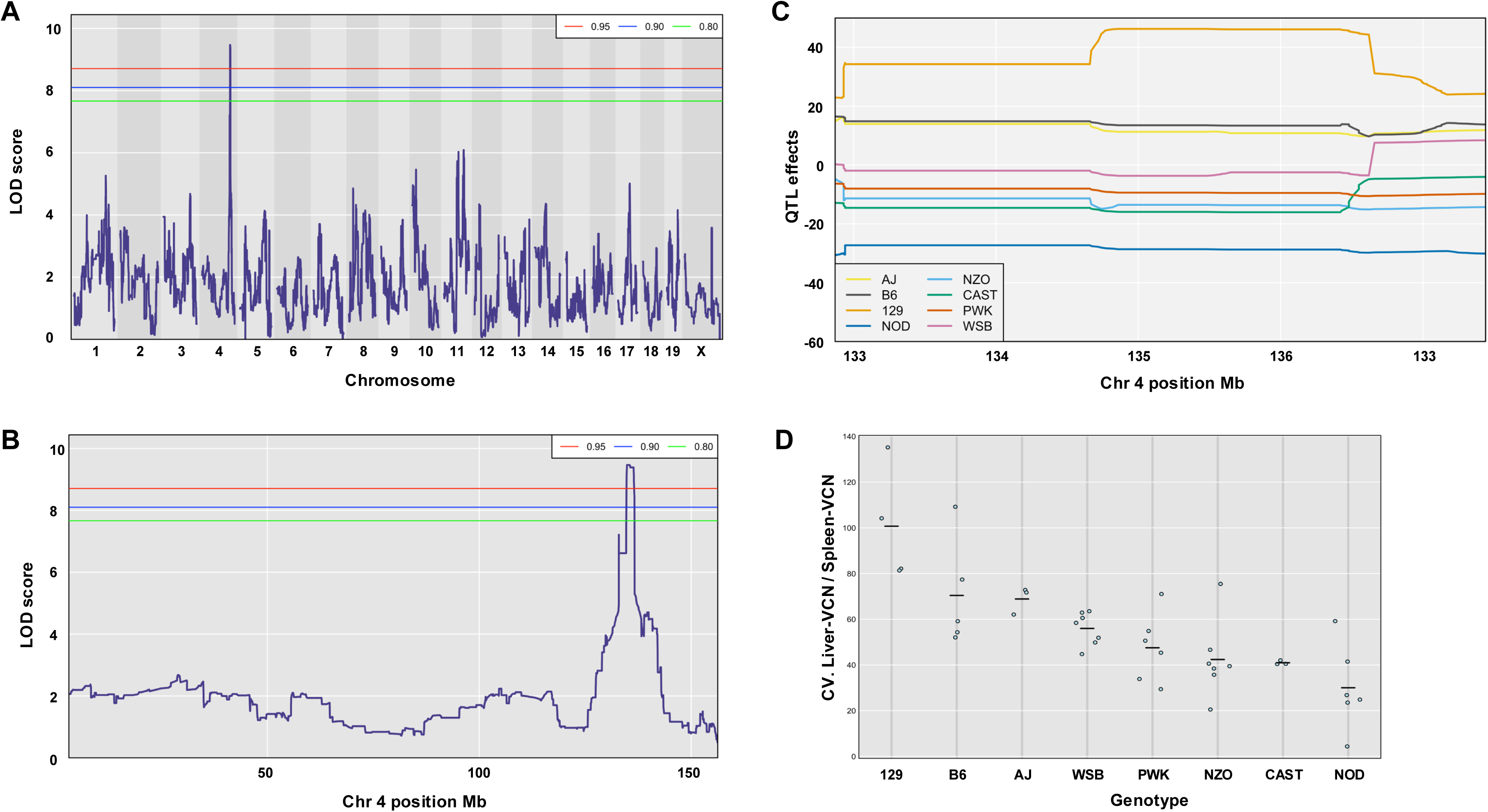
Host genetic background effects on intra strain variation in the ratio of liver and spleen VCN. CV (SD/mean) of the ratios of liver and spleen VCN in each CC mouse strain was employed to identify genomic loci associated with intra strain variation in the above phenotype. A. shows the LOD scores across all chromosomes. The horizontal red, blue, and green lines indicate the 5%, 10%, and 20% genome-wide significance threshold, respectively, based on a permutation test. B. The LOD score on chromosome 4. C. The estimated QTL effects of eight haplotypes among markers near the peak of QTL. D. Dot plot showing the CV of the ratio of liver and spleen VCN in CC mouse stains carrying one of the 8 founder alleles at the the above QTL on chromosome 4. Note that 4 mouse strains carrying the 129 haplotype exhibit the highest CVs.

**Figure 19.**
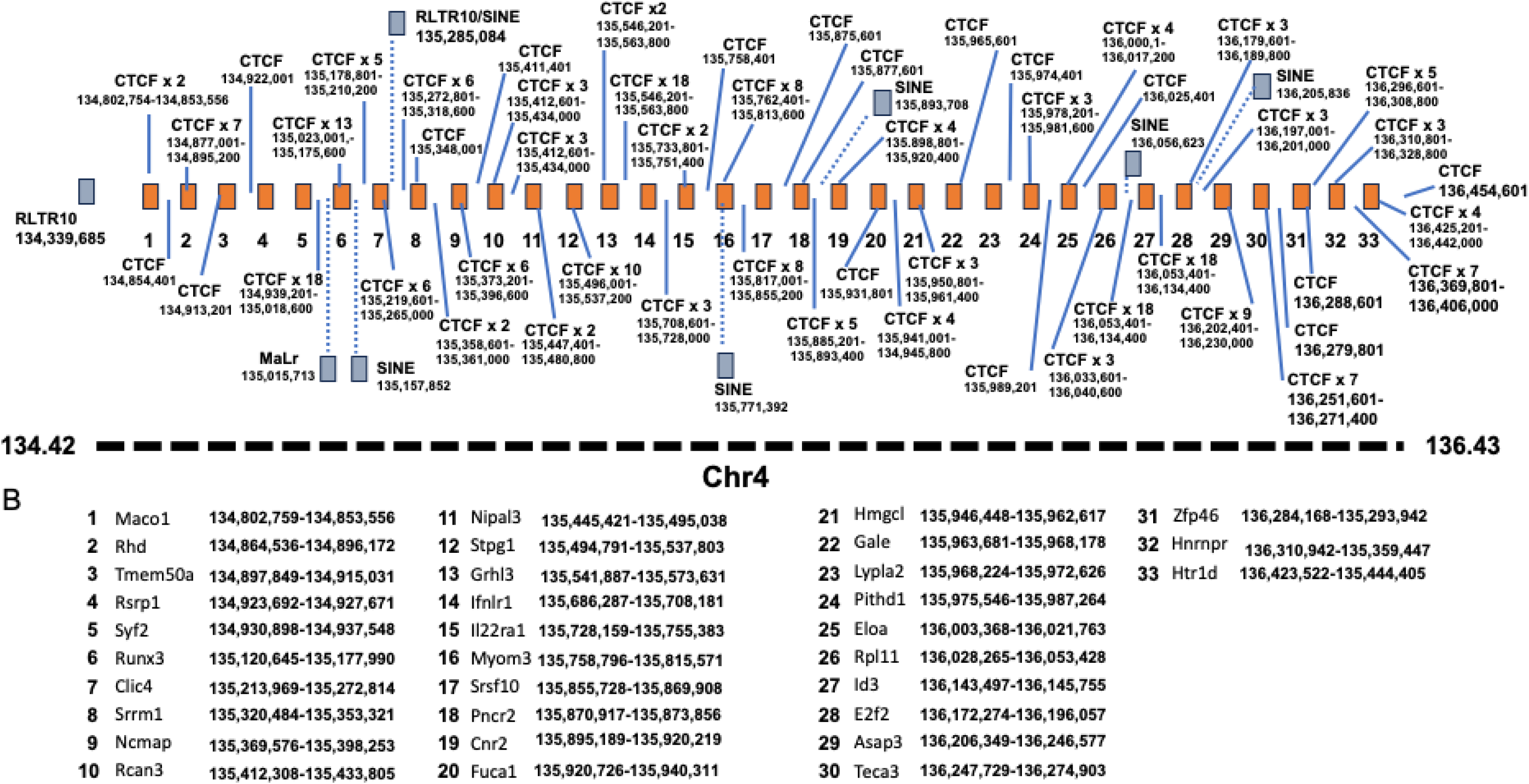
A physical map of a QTL in mouse chromosome 4, which associates with the ratios of liver and spleen VCNs. A. Depiction of the above QTL located between Mbs 134.42 and 136.43 in chromosome 4. Numbered red boxes show the genes comprised in the above QTL. Blue boxes show the various transposable elements and their location in chromosome 4 (determined by Nellaker *et al.* 2012 (43) and converted to Ensmbel GRCm38.6 by https://genome.ucsc.edu/cgi-bin/hgLiftOver). CTCF target sequences and their location in chromosome 4 are shown (determined by Ensmbel GRCm38.6). B. Names and locations of the respective numbered red boxes shown in the above figure 19 A.

### QTL association with CV of hepatic luciferase expression at week 3-PVA

A second QTL with LOD score of 8.92 and p-value of 0.030 was associated with CV (strain-specific intra-strain variation) of hepatic luciferase expression levels at week 3-PVA (Figures 20 A-D and Table 4). The QTL is positioned in 50.19 Mb of chromosome 7. Its width is 9.28 Mb, and it comprises 372 genes and 134 pseudogenes. Its percentage of contribution (heritability) to the phenotype is 63.26%. As shown in figures 20 C-D and table 4, CC mouse strains (CC002, CC003, CC033, CC058), which are composed of the PWK allele in the above QTL are associated with the highest CV.

**Figure 20.**
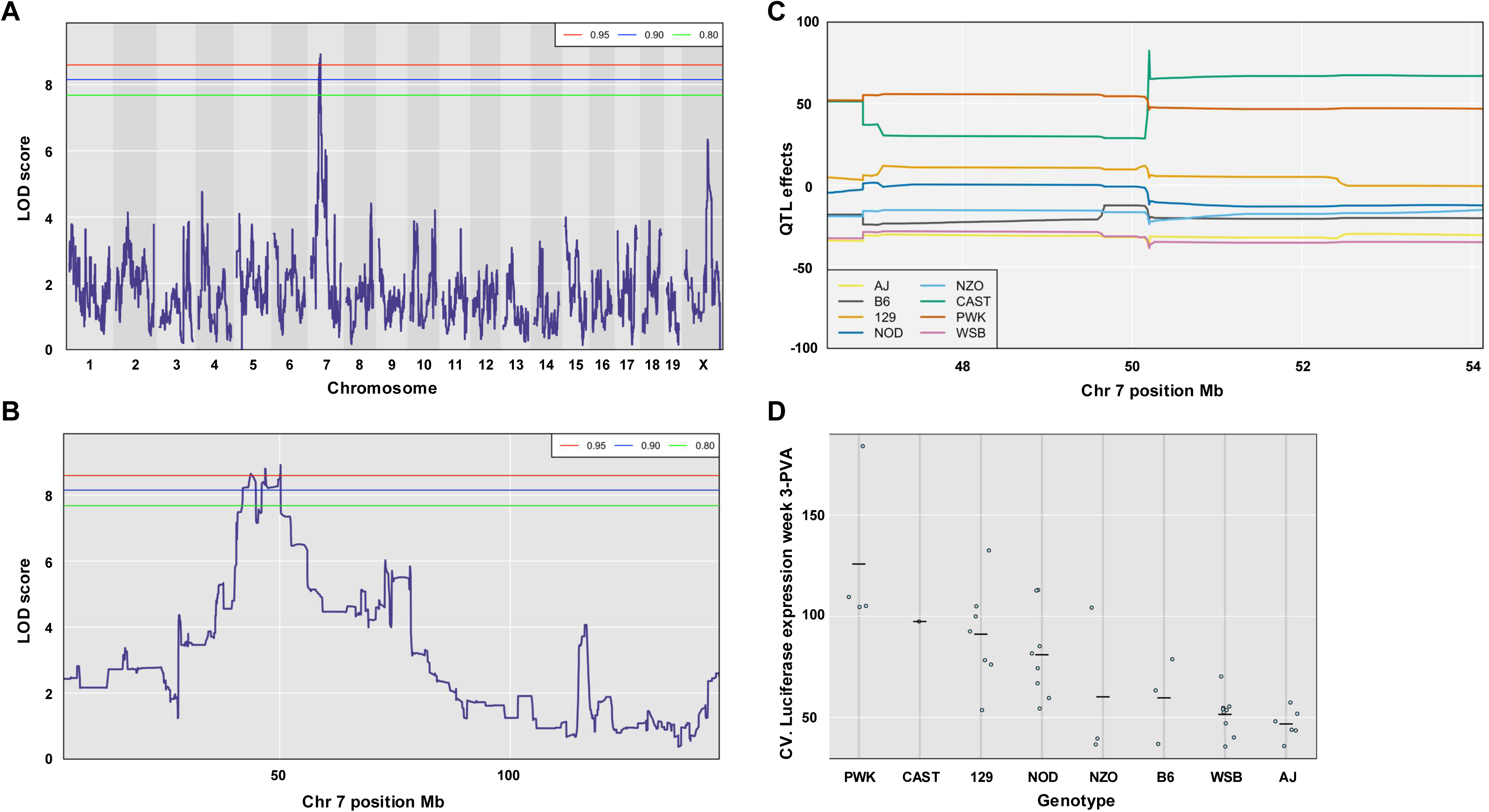
Host genetic background effects on intra strain variation in hepatic luciferase expression at week 3-PVA. CV (SD/mean) of hepatic luciferase expression at week 3-PVA in each CC mouse strain was employed to identify genomic loci associated with intra strain variation. A. The LOD scores across all chromosomes. The horizontal red, blue, and green lines indicate the 5%, 10%, and 20% genome-wide significance threshold, respectively, based on a permutation test. B. The LOD score on chromosome 7. C. The estimated QTL effects of eight haplotypes among markers near the peak of QTL. D. Dot plot showing the CV hepatic luciferase expression at week 3-PVA in CC mouse stains carrying one of the 8 founder alleles at the the above QTL on chromosome 7. Note that 4 mouse strains carrying the PWK haplotype exhibit the highest CVs.

### Identification of a QTL associated with the increase in hepatic luciferase expression between weeks 1- and 3-PVA

The kinetics of hepatic transgene expression following nuclear import of lentiviral vectors is one of the factors that determine the outcomes of gene therapy protocols. To identify genomic loci, which associate with changes in hepatic transgene expression from lentiviral vectors, differences between hepatic luciferase expression levels at weeks 1- and 3-PVA were determined. To minimize intrinsic effects of transgene expression levels on the ability to associate the kinetics of transgene expression and specific QTLs, the above differences in expression levels at weeks 1- and 3-PVA were normalized by dividing the above differences in luciferase expression with the levels of luciferase expression at week 1-PVA. As shown in figures 21 A-D and table 4, QTL analysis identified a single QTL, with LOD score of 8.37 and p-value of 0.04. It was associated with an increase in transgene expression between week 1 and week 3-PVA. The QTL is positioned at 192.36 Mb of Chromosome 1. It encompasses 1.35 Mb, and the top SNP in QTL regions contributes 60.92% to the above phenotype. As shown in figures 21 C-D, 5 CC mouse strains (CC003, CC012, CC019, CC061, CC063), which are composed of the 129s1/SvImJ allele in the above QTL are associated with the highest increase in transgene expression between week 1- and week 3-PVA. Importantly, the above QTL interval comprises 34 genes and 4 pseudogenes including 5 protein encoding sequences, which potentially affect the kinetics of transgene expression between week 1- and week 3-PVA. These genes include: A) *SERTAD4* (involved in cell cycle progression and chromatin remodeling) ^48,49^, B) *IRF6* (regulator of development and the innate immune response) ^50-53^, C) *TRAF3IP* (regulator of the innate immune response) ^54,55^ *and* D) *UTP25/DIEFX* (regulator of development, ribosome biogenesis and tumor development) ^56-58,59^. Importantly, a recent study ^60^ reported on chromatin-based interactions between genetic variants of the UTP25 promoter, and the *TRAF3IP3* and *IRF6* genes in immune cells which are involved in an autoimmune disorder.

**Figure 21.**
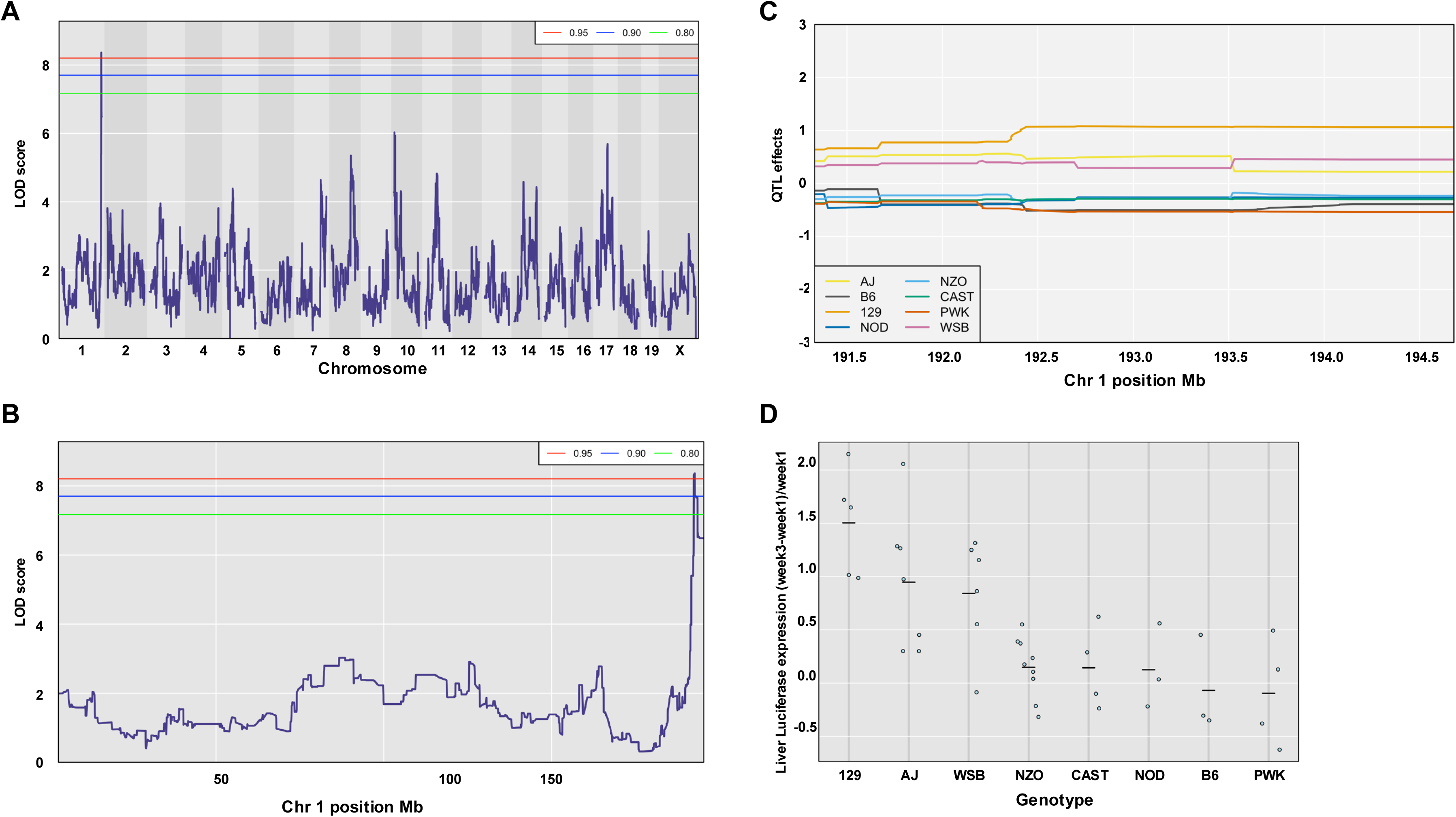
Characterization of host genetic background effects on the kinetics of hepatic luciferase expression between weeks 1 and 3-PVA. To this end the differences in hepatic luciferase expression between weeks 1 and 3-PVA were normalized (divided) by the level of hepatic luciferase expression at week 1-PVA. A. The LOD scores across all chromosomes. The horizontal red, blue, and green lines indicate the 5%, 10%, and 20% genome-wide significance threshold, respectively, based on a permutation test. B. The LOD score on chromosome 1. C. The estimated QTL effects of eight haplotypes among markers near the peak of QTL. D. Dot plot showing the means of normalized differences in hepatic luciferase between weeks 1 and 3-PVA in CC mouse stains carrying one of the 8 founder haplotypes at the the above QTL on chromosome 1. Note that 5 mouse strains carrying the 129 haplotype exhibit the highest increase in hepatic luciferase expression between weeks 1 and 3-PVA.

## Discussion

Empirically tailoring conventional therapeutic regimens to an individual patient, only partially compensate for our limited ability to accurately predict the safety and efficacy of therapeutic protocols for an individual patient. Furthermore, this methodology cannot be applied to gene therapy protocols, which are premised on a single administration procedure.

Variability in the outcome of viral vector-based gene therapy clinical trials can be attributed to patient-specific expression profiles of viral dependence and viral restriction factors. An increasing number of host factors have been identified as HIV dependence factors (HDFs), which are necessary for efficient progression through the early and late stages of the viral life cycle.^17-21^ HIV restriction factors (HRFs) are the host’s first line of antiviral defense, which primarily target the early steps of the viral life cycle. Although, already present in host cells and ready to inhibit viral replication at the time of infection, HRFs also serve as inducers of the innate immune response, which in turn further enhance HRFs expression.^61,62^

In contrast to the multistep HIV-1 life cycle, the process of gene-delivery by non-replicating HIV-1 vectors is a single round of infection event, which starts at vector attachment to the target cells and is completed with the initiation of transgene expression. In contrast to a productive HIV life cycle, which results in the release of infectious viral particles, successful transduction by lentiviral vectors leads to long-term transgene expression and survival of vector-transduced cells.

For more than a decade various approaches have been employed by different groups to identify host factors affecting HIV-1 infection *in vitro.* ^63-65^ Interestingly, results from these studies demonstrated only a limited overlap. By default, the above studies could not identify candidate genes, which contribute to the host *in vivo* environment and to the adaptive immune response to HIV-1 infection, which affect long-term expression of viral genes (e.g. HIV latency). Furthermore, *in vitro* screening methodologies which are premised on supraphysiological expression levels of transgenes or highly efficient knockdowns of host gene expression are probably not compatible with normal human physiology. Consequently, small molecules that efficiently target the above *in vitro* identified HDFs and HRFs are likely to inadvertently induce cytotoxicity, which limits their therapeutic value.

Small animal models using a single mouse strain are the premise of most safety and efficacy preclinical studies. The applicability of results obtained in mouse models to human translational research could be limited and probably mouse strain specific. Importantly, recent mouse models employing the EcoHIV chimeric HIV virus accurately emulated complex phenotypes observed in HIV infected patients including HIV latency and HIV induced neurocognitive impairment.^66,67^ Per contra, a cardinal study aimed at modeling the development of cellular immune response to AAV vector-transduced hepatocytes in gene therapy treated patients, did not exhibit characteristics of cellular immune response to AAV vector transduced hepatocytes in immunologically sensitized mice.^68^ These findings were not in line with the outcomes of an earlier gene therapy clinical trial, in which hemophilia B patients treated with factor IX expressing AAV2 vectors exhibited decline in therapeutic factor IX levels and an increase in hepatic transaminases.^13^ Recently reported discrepancies between stable long-term correction of factor VIII deficiency in preclinical studies of AAV vector-based gene therapy for hemophilia A and human clinical studies underscore the significance of this phenomenon.^69^ The discrepancies between the above animal model and the outcomes of the AAV vector-based human clinical trial were attributed to several mechanisms including differences in the function of mouse and human CD8 cells as well as differences in mouse and human AAV capsid processing and antigenic presentation.^68^ We raise the possibility that mouse strain-specific traits could also contribute to the discrepancies between human clinical trials and their respective mouse preclinical studies, which are premised on a single mouse strain. Importantly, in the current study we employed 42 genetically defined mouse-strains to characterize the effects of the host genetic background on multiples parameters of hepatic gene delivery by lentiviral vectors. Interestingly, a comprehensive study by Uchil et al. demonstrated differences in the effects of mouse and human Trims on HIV transduction efficiency of human and mouse cells. These finding suggest that additional mechanism contributes the discrepancies between preclinical and clinical studies.^70^

Throughout the study we observed significant differences in hepatic transgene expression among the 41 CC mouse strains (e.g. more than 100-fold difference in hepatic luciferase expression in CC057 and CC061). Importantly, wide phenotypic range across the CC mouse strain was common to all analyzed traits including hepatic and splenic VCN, and vector specific activity. These findings indicate that the mouse genetic background and by extrapolation the genetic background of human patients determine the efficacy and safety of preclinical and therapeutic lentiviral vector-based gene delivery protocols, respectively. We assert that significant variability in the outcomes of currently employed gene therapy protocols should be expected. Also, it appears that unexpected outcomes of gene therapy clinical trials cannot be predicted by currently employed murine-based preclinical studies.^68^ We assert that in order to compensate for the limited ability to predict the outcomes of a gene therapy protocol in an individual patient, it is reasonable to consider the use of escalating dose-based protocols.

In most CC mouse strains, periodic analysis of hepatic luciferase expression demonstrated stable or an increase in transgene expression between weeks 1 and 24-PVA. However, two CC mouse strains (CC021 and CC024) exhibited continuous decrease in hepatic luciferase expression. The mean hepatic VCN of the CC021 strain (0.41 vector genomes per cell) was higher than the VCN of 24 CC mouse strains. Based on these findings we theorized that epigenetic silencing of lentiviral vectors carrying hepatic specific promoter (hAAT) contributed to this phenomenon. This notion was supported by the fact that vector specific activity (SA) in CC021 hepatocyte was the lowest among the studied CC mouse strains. In contrast, CC024 mouse strain hepatocytes exhibited the lowest mean VCN of 0.045 and an average SA, which suggested that either luciferase cytotoxicity or the host cellular immune response to luciferase expressing hepatocytes were involved with the decline in hepatic luciferase expression between weeks 1 and 24-PVA.^42^ The continuous decline in hepatic luciferase expression in CC021 and CC024 was mediated by two separate mechanisms. It took place in the context of healthy animals, which were transduced with integrating, self-inactivating (SIN) vectors carrying a liver specific promoter. These characteristics suggest that the above mechanisms are probably part of a general host antiviral response, which is independent of the HIV LTR sequence, the vector integration site, and the nature of the vector’s internal promoter. These findings open new directions in identifying novel host factors, which affect long-term expression from lentiviral vectors and could be involved in the establishment of latent HIV reservoirs. Furthermore, since the above phenomena were characterized in healthy mice, it is reasonable to assume that mild non-toxic alterations in expression/function of the above newly identified host factors could be sufficient to prevent the decline in hepatic transgene expression from lentiviral vectors and to treat the latent HIV reservoirs with minimal adverse effects. Interestingly, to a certain degree the continuous decrease in luciferase in CC024 and CC021 is in line with the decrease in factor FVIII activity in hemophilia A patients treated with high doses of AAV vectors carrying factor VIII.^69^

The notion that preclinical mouse studies can be employed to characterize complex, multifactorial pathologic traits in humans was the impetus to the development of the CC recombinant inbred mouse strains panel.^71^ As described above, 8 mouse strain were subjected to a carefully planned intercrossing breeding protocol, which generated novel, genotypically characterized mouse strains. The novel CC panel of mouse strains has been successfully employed in an increasing number of studies aimed at understanding the role of the genetic background in the development and outcomes of various pathologic processes. These include bacterial, fungal and viral infections, ^24,25,72-77^ metabolic disorders, ^78^ inflammation ^26^, cancer ^79-81^ neurological and immunologic disorders.^82,83^ Here we employed the valuable CC mouse strains panel to identify host factors that affect a therapeutic procedure.

The CC panel was designed to genetically characterize complex traits including the identification of allelic combinations that quantitatively affect complex traits. It was estimated that 1000 CC mouse strains would be required to meet the above objectives.^71^ At this time, the merely 70 CC mouse strains that are available, cannot provide the statistical power to accurately characterize highly complex traits. Although some of the above studies successfully associated QTLs (and at times specific host factors) with specific phenotypes, the statistical power of the CC mouse panel was not sufficient to identify specific combinations of normal alleles that quantitively affected complex traits. Notwithstanding the statistical limitations of the current CC panel, it is a most powerful research tool that provides novel insights that expand our understanding of biological pathways. The wide genetic variability within the CC panel accurately outlines the spectrum of strain-specific phenotypes and identifies specific CC mouse strains, which exhibit the broadest strain-specific phenotypic divergence. This critical attribute facilitates further genetic characterization of the above phenotypes by F2-crossing studies (employing 2 extreme strains).

The current study aimed at identifying host factors that determine the ultimate outcome of hepatic gene delivery by lentiviral vector, which was defined here as the mean of hepatic luciferase expression level at week 24-PVA. Analysis of mouse strain genotypes and hepatic luciferase expression failed to identify a QTL that associates with hepatic luciferase expression at week 24-PVA. As outlined earlier, successful hepatic transduction, which results in long-term transgene expression is a multistep process. Each step in this process is affected by multiple HRFs and HDFs, which are encoded by 8 different alleles. Unless a highly potent allele is comprised in the genetic pool of the 8 CC founders, it is likely that the statistical power of a 41 CC mouse strain-based genetic study will not be sufficient to associate a QTL with hepatic luciferase expression at week 24-PVA.

To circumvent the above statistical power limitations in the current panel of CC mouse strains, we sought to narrow the complexity of the studied phenotypes. To this end, we defined the relative differences in hepatic luciferase expression between weeks 1 and 3-PVA (normalized by the level of luciferase expression at week 1-PVA (Luc-W3 - Luc-W1 / Luc-W1)) as a new trait, which exhibited the highest correlation with hepatic luciferase expression at week 24-PVA. This approach is premised on the notion that in contrast to the multitude of host factors that affect multiple steps in the process of hepatic lentiviral transduction, a smaller number of host factors are involved in the mechanism that affects the kinetics of hepatic transgene expression between weeks 1 and 3-PVA. We theorized that the narrowed trait complexity would increase the likelihood of associating a QTL with the phenotype of interest. Indeed, a QTL in chromosome 1 was associated with an increase in hepatic luciferase between weeks 1 and 3-PVA. The 129s1/SvImJ founder allele in the above QTL was shared by 5 CC mouse strains, which exhibited the highest increase in hepatic luciferase expression between weeks 1 and 3-PVA. Within the above QTL we identified 4 candidate genes, which can potentially contribute (individually or synergistically) to the studied trait. These genes include *SERTAD4, IRF6, TRAF3IP3 and UTP25,* which are involved in embryo development, cancer, innate immune response and epigenetic processes. ^49,50,52,54^ Interestingly, the impact of *TRAF3IP3* on interferon pathways^54^ could be in line with earlier reports demonstrating interferon effects on hepatic lentiviral vector transduction.^42^ The relative proximity of the above genes in chromosome 1 raises the possibility that their expression is coregulated. Interestingly, findings from a recent study indicated that genetic variants in either the *UTP25* or the *IRF6* promoters affect their interactions with either *IRF6* and *TRAF3IP3* or *UTP25* and *TRAF3IP3*, respectively.^60^ These interactions had different effects on the expression level of the above genes. Further research is required to determine the effects of each of the above candidate genes on lentiviral transduction.

During this study, we observed phenotypic variability among mice of the same strain (Isogenic-phenotypic discordance). The level of the observed isogenic discordance, (quantified as coefficient variation) was mouse-strain and phenotype dependent. Phenotypic discordance between monozygotic twins has been associated with several human diseases. ^84^ The phenomenon of isogenic discordance has been mechanistically studied in different animal models.^85,86^ One of the first murine models studying isogenic discordance, was premised on mice carrying the *agouti viable yellow (A^vy^)* allele.^87,88^ *A^vy^* mice exhibit various fur colors ranging from agouti to complete yellow, which are secondary to the expression level of the agouty-protein (Yellow and agouti fur colors are associated with high and low levels of the agouti protein, respectively). This phenomenon is attributed to germline insertion of an intracisternal A-type particle (IAP) transposon upstream to the agouti protein coding sequence. Consequently, transcriptional regulation of the *A^vy^* allele is mediated by the IAP LTR, whose methylation inversely correlates with its transcriptional activity and varies among isogenic *A^vy^* mice. Isogenic alleles which demonstrate variable epigenetic profiles and consequently different transcriptional activity are termed metastable epialleles.^89-91^

The unique DNA methylation patterns of IAP LTRs are established during gametogenesis and early embryogenesis which involve epigenetic reprograming processes including 2 waves of genome-wide DNA de-methylation (in primordial germ cells and preimplantation embryos) followed by de-novo methylation.^92-95^ Partial methylation of IAP LTRs in the preimplantation embryo and during gametogenesis indicate that the IAP LTRs are relatively resistant to the above genome wide demethylation processes.^94-96^ Early publications suggested that DNA hypermethylation, which represses expression of IAPs and other retrotransposons (especially during gametogenesis) is part of a host defense mechanism that minimizes genomic instability due to mutagenic germ line insertions of retro-elements.^97,98^ On a cellular level (which is mostly mouse-strain independent), the phenomena of genomic imprinting and chromosome X inactivation in female cells result in allele-specific gene expression. Interestingly, an epigenetic modifier that affect expression of imprinted genes also associates with IAP genomes.^47^ Furthermore, the fur color of *A^vy^*mice is affected by the maternal phenotype, which is a genomic imprinting characteristic.^44,88,90^

Most IAP genomes are transcriptionally silenced by suppressive histone modifications and DNA methylation.^89,94,96,99^ However, specific IAP proviruses show variable levels of DNA methylation and maintain variable transcriptional activity in a locus specific manner in isogenic mice. In IAP-comprising metastable alleles, the process that determines the methylation status of a specific IAP is stochastic. However, the overall methylation status of a metastable allele in specific mouse strains is probabilistic and affected by several variables including the IAP sequence, the IAP integration site and the mouse strain genetic background ^100-102^ which determines the strain specific expression profile of multiple epigenetic modifiers.^103-105^ Importantly KRAB-ZFPs which have a major mechanistic role in epigenetic repression of IAP transcription have been evolved to target strain-specific IAP sequences.^89,106^ Several studies reported on environmental effects on the epigenetics and gene expression of metastable epialleles. These include nutritional alteration, exposure to ionizing radiation, heavy metals, ethanol and bisphenol A (BPA).^107-111^ However, later studies could not fully support some of the above earlier reports.^112,113^

Notwithstanding the central role of IAP sequences in the metastable epiallelic phenomena, the Whitelaw team described variegation of transgene expression from tandems of expression cassettes lacking IAP sequences.^114,115^ The above transgenic construct contained the green fluorescence protein (GFP) open reading frame under the control of the human α-globin promoter. Variegation levels of erythrocyte GFP expression were mouse clone specific. However, intra clone variability of variegation level (between mice within a specific mouse clone) has not been reported. Furthermore, using a germ line chemical mutation procedure, the Whitelaw team identified secondary epigenetic modifiers which either enhanced or suppressed clonal variegation and altered expression of bona-fide metastable epialleles in *A^vy^* mice.^114^ These findings suggest that unique *cis* elements such as IAP sequences and imprinting control elements (ICE) are the primary initiators of the metastable epigenetic phenomenon, which is premised on local epigenetic permutations at the time of genome wide epigenetic reprograming processes during either gametogenesis, preimplantation embryo and probably in some terminal differentiation processes.^116-119^ It is likely that association of specific trans elements such as KRAB-ZFPs with the above primary *cis* elements is an early (and probably a required) event in the establishment of metastable epiallelic hubs.^47,120,121^ This early event is followed by interactions with the above secondary epigenetic modifiers whose levels affect the range of metastable allele expression (and DNA methylation) among isogenic mice.^114^ Per contra genome-wide epigenetic reprograming processes during cellular differentiation ^116-119^ can potentially results in variegation of transgene gene expression whose range is also determined by the above epigenetic modifiers described by Blewitt *et al*. ^114^ It is possible that during genome-wide epigenetic changes, (typical of gametogenesis, early embryo development and cellular differentiation) a balance between two pathways including cellular differentiation (premised on DNA demethylation and is CTCF dependent) and repression of transposable element (premised on epigenetic transcriptional repression requiring KRAB-ZFPs) results in isogenic phenotypic discordance. In this study we observed phenotypic-isogenic discordance in several parameters of hepatic transduction by lentiviral vectors. In contrast to earlier transgenic mouse models, the lentiviral vector genomes in this study did not pass through gametogenesis or early preimplantation embryonic processes. Furthermore, the integration site profile of lentiviral vectors is semi-random,^122^ thus the likelihood that significant number of lentiviral vectors integrated in proximity to IAP sequences is low. Note that the self-inactivating (SIN) lentiviral vectors in this study were devoid of the HIV U3 region (which comprises the parental HIV enhancer promoter sequences). Although, endogenous lentiviral sequences were identified in rodent and primate genomes, endogenous murine lentiviral genomes have not been detected.^123-126^ These facts minimize the likelihood that lentiviral vector sequence-directed KRAB-ZFPs contributed to the intrastrain phenotypic variations observed in this study. However, we cannot entirely rule out the possibility that KRAB-ZFPs directed to murine retroelements cross reacted with the lentiviral vector genomes. Since mouse hepatic transduction by lentiviral vectors is highly regulated by HRFs and HDFs, which also regulate the life cycle of murine endogenous and exogenous retroviruses, we raise the possibility that variation in the expression levels of the above host factors in isogenic mice probably contributed to the intrastrain phenotypic variations, which were characterized in this study. The large number of the host factors involved in the various steps of lentiviral vector hepatic transduction and the number of mouse strain-specific transposable elements (TE) in the mouse genome support the above notion.^45^ The wide range of mouse-strain specific CVs of multiple lentiviral vector transduction traits raised the possibility that the phenomenon of genetic and phenotypic discordance is not rare. To characterize the role of the host genetics in the mechanism of phenotypic variability in isogenic mice, a LOD score analysis was employed to identify two QTLs. A QTL of 9.28Mb (comprising 372 and 134 genes and pseudogenes, respectively) in chromosome 7 was associated with CV of hepatic luciferase expression at week 3-PVA. A second and significantly narrower QTL of 1.61Mb in chromosomes 4 was associated with the CV level in the ratio of liver and spleen VCNs. Specifically, increased CV levels were associated with the 129s1/SvImJ allele in the above QTL in three CC mouse strains. Premised on an earlier study, ^45^ further QTL characterization identified eight 129s1/SvImJ-specific insertions of retroelements including 5 SINEs, 2 RLTR10 and a single MaLr insertion, which was located within a cluster of 8 CTCF target sequences. The central role of CTCF and its interactions with transposable elements in the phenomena of phenotypic variation in isogenic mice and in genomic imprinting, as well as in strain-specific variation in gene expression levels has been reported earlier. Although, the 129s1/SvImJ allele in the above QTL was associated with the highest CV it does not contain strain-specific ^43-46^ IAP insertions, whose variable methylation (VM) status has been mechanistically associated with the phenomenon of metastable epialleles.^43,87,88^ Importantly, although not common, VM of non-IAP retroelements have been reported by recent studies.^43,44^ However, their effect on intrastrain variation in gene expression has not been documented. The studied phenotype (CV of splenic VCN / hepatic VCN) was premised on lentiviral VCN in 2 tissues. There is a possibility that expression of a host factor affecting VCN exhibits intrastrain variability in a tissue specific manner. This phenomenon was described in an earlier report describing tissue specific variable methylation of IAP sequences (tsVM-IAP) in isogenic animals.^43^ However, tsVM of non-IAP retroelements and its effect on gene expression has not been reported. Note that the human genome is devoid of IAP sequences. Thus, the phenomenon of metastable epialleles in humans^91,127^ is likely to involve non-IAP transposons.

The ultimate dependency of viruses on host physiologic and metabolic pathways, which are probably modulated by the host circadian rhythm ^128,129^ conjure up the notion that host sleep patterns affect lentiviral vector transduction efficiency. An earlier study demonstrating major circadian rhythm effects on influenza A virus (IAV) and herpes simplex virus 1(HSV-1) infection *in vivo* ^130^ supported the above notion and was the main impetus to correlate sleep patterns of CC mouse strains with overall lentiviral vector transduction efficiency as determined by hepatic luciferase expression levels at week 24-PVA. This study describes for the first-time moderate correlation between behavioral/sleep patterns and *in vivo* lentiviral vector transduction efficiency. Our findings suggest that the host sleep pattern is one of various host physiologic pathways that determine the outcomes of lentiviral vector transduction. Furthermore, as outlined by Sulli et. al.,^128^ the circadian rhythm has relatively modest effects on gene expression. Two mechanisms can separately or in combination contribute to this phenomenon. One mechanism requires that host factors affecting sleep patterns are also involved in the multistep process of hepatic transduction. However, there is also a possibility that maternal behavior/sleep pattern from the time of gametogenesis to the progenies preweaning period can epigenetically induce lifelong alterations in gene expression, which affect permissiveness to lentiviral vector transduction.^86,131-133^ Note that notwithstanding the circadian rhythm effects on multiple physiologic pathways and the outcomes of viral infections,^128-130,134,135^ mouse strains with altered melatonin function are the premise of animal models in basic research and preclinical studies.^136,137^ Furthermore, animal care in most academic animal facilities is done during the mouse daylight/sleep period. The animal care-induced sleep disturbances can be considered as an environmental factor, whose effects on mouse sleep patterns are mouse-strain dependent which can contribute to the phenomenon of intrastrain phenotypic differences.

## Conclusions

In this study 41 genetically characterized CC mouse strains were employed to investigate the effects of the host genetic background on hepatic transduction by lentiviral vectors.

The study demonstrated a wide range of mouse strain specific differences in various aspects/traits of lentiviral vector hepatic gene delivery. These major host genetic background effects on lentiviral vector transduction *in vivo* suggest that novel multiple mouse strains-based approaches of evaluating the efficacy and safety of viral vector-based gene delivery in preclinical studies should be established. We believe that the new approaches would narrow current discrepancies between the outcomes of preclinical to clinical studies and will be highly cost-effective.

The genetic diversity of human patients, which is not narrower than the genetic diversity of 41 CC mouse strains explains the wide range of patient-specific outcomes of viral vector-based gene therapy protocols. It appears that the current phase I to III system of clinical trials may not fully address the diversity of human genetics. We assert that accumulating or escalating dose-based protocols of viral vector administration should be considered for gene therapy protocols for non-fatal diseases.

Using CV of various gene delivery phenotypes associated 2-QTLs with intrastrain phenotypic variations. These findings suggest that the phenomenon of phenotypic isogenic discordance in mice is not rare and can be further characterized by using large number of genetically defined mouse strains.

In line with earlier reports,^130^ this study correlated for the first time the efficiency of lentiviral transduction *in vivo* with strain-specific sleep patterns. The findings of this study raised the possibility that routine mouse care procedures in most animal facilities induce sleep disturbances, which have strain specific effects on HIV-based vector transduction.

## Acknowledgements

The following reagents were obtained through the National Institutes of Health (NIH) AIDS Research and Reference Reagent Program, Division of AIDS, the National Institute of Allergy and Infectious Diseases: the HIV-1 p24 monoclonal antibody (183-H12-5C) from Bruce Chesebro and Kathy Wehrly. The study was supported by NIH grants R01-HL128119 (to TK and PH) and R01-HL155986 (to TK, FZ, YH, GD and TW). In memory of all the women in the black shirt. In honor of B’Tselem. Genesis 1:27.

